# Modeling Cell-Specific Dynamics and Regulation of the Common Gamma Chain Cytokines

**DOI:** 10.1101/778894

**Authors:** Ali M. Farhat, Adam C. Weiner, Cori Posner, Zoe S. Kim, Brian Orcutt-Jahns, Scott M. Carlson, Aaron S. Meyer

## Abstract

Many receptor families exhibit both pleiotropy and redundancy in their regulation, with multiple ligands, receptors, and responding cell populations. Any intervention, therefore, has multiple effects, confounding intuition about how to precisely manipulate signaling for therapeutic purposes. The common γ-chain cytokine receptor dimerizes with complexes of the cytokines interleukin (IL)-2, IL-4, IL-7, IL-9, IL-15, and IL-21 and their corresponding “private” receptors. These cytokines have existing uses and future potential as immune therapies due to their ability to regulate the abundance and function of specific immune cell populations. However, engineering cell specificity into a therapy is confounded by the complexity of the family across responsive cell types. Here, we build a binding-reaction model for the ligand-receptor interactions of common γ-chain cytokines enabling quantitative predictions of response. We show that accounting for receptor-ligand trafficking is essential to accurately model cell response. This model accurately predicts ligand response across a wide panel of cell types under diverse experimental designs. Further, we can predict the effect and specificity of natural or engineered ligands across cell types. We then show that tensor factorization is a uniquely powerful tool to visualize changes in the input-output behavior of the family across time, cell types, ligands, and concentration. In total, these results present a more accurate model of ligand response validated across a panel of immune cell types, and demonstrate an approach for generating interpretable guidelines to manipulate the cell type-specific targeting of engineered ligands. These techniques will in turn help to study and therapeutically manipulate many other complex receptor-ligand families.

**Summary points:**

- A dynamical model of the γ-chain cytokines accurately models responses to IL-2, IL-15, IL-4, and IL-7.
- Receptor trafficking is necessary for capturing ligand response.
- Tensor factorization maps responses across cell populations, receptors, cytokines, and dynamics to visualize cytokine specificity.
- An activation model coupled with tensor factorization provides design specifications for engineering cell-specific responses.

## Introduction

Cytokines are cell signaling proteins responsible for cellular communication within the immune system. The common γ-chain (γ_c_) receptor cytokines, including interleukin (IL)-2, 4, 7, 9, 15, and 21, are integral for modulating both innate and adaptive immune responses. As such, they have existing uses and future potential as immune therapies.^1,2^ Each ligand binds to its specific private receptors before interacting with the common γ_c_ receptor to induce signaling.^3^ γ_c_ receptor signaling induces lymphoproliferation, offering a mechanism for selectively expanding or repressing immune cell types.^4,5^ Consequently, loss-of-function or reduced activity mutations in the γ_c_ receptor can cause severe combined immunodeficiency (SCID) due to insufficient T and NK cell maturation.^6^ Deletion or inactivating mutations in IL-2 or its private receptors leads to more selective effects, including diminished regulatory T cell (T_reg_) proliferation and loss of self-tolerance.^7–9^ Deficiency in the IL-2 receptor IL-2Rα also causes hyperproliferation in CD8+ T cells, but diminished antigen response.^10^ These examples show how γ_c_ receptor cytokines coordinate a dynamic balance of immune cell abundance and function.

The γ_c_ cytokines’ ability to regulate lymphocytes can impact both solid and hematological tumors.^11^ IL-2 is an approved, effective therapy for metastatic melanoma, and the antitumor effects of IL-2 and IL-15 have been explored in combination with other treatments.^12,13^ Nonetheless, understanding these cytokines’ regulation is stymied by their complex binding and activation mechanism.^3^ Any intervention imparts effects across multiple distinct cell populations, with each population having a unique response defined by its receptor expression.^14,15^ These cytokines’ potency is largely limited by the severe toxicities, such as deadly vascular leakage with IL-2.^16^ Finally, IL-2 and IL-15 are rapidly cleared renally and by receptor-mediated endocytosis, limiting their half-life *in vivo*.^17–19^

To address the limitations of natural ligands, engineered proteins have been produced with potentially beneficial properties.^2^ The most common approach has been to develop mutant ligands by modulating the binding kinetics for specific receptors.^20,21^ For example, mutant IL-2 forms with a higher binding affinity for IL-2Rβ, or reduced binding to IL-2Rα, induce greater cytotoxic T cell proliferation, antitumor responses, and proportionally less T_reg_ expansion.^12,22^ This behavior can be understood through IL-2’s typical mode of action, in which Tregs are sensitized to IL-2 by expression of IL-2Rα.^14^ Bypassing this sensitization mechanism thus shifts cell-specificity.^22^ Conversely, mutants skewed toward IL-2Rα over IL-2Rβ binding selectively expand T_reg_ populations, over cytotoxic T cells and NK cells, compared to native IL-2.^23,24^

The therapeutic potential and complexity of this family make computational models especially valuable for rational engineering. Early attempts at mathematically modeling the synergy between IL-2 and IL-4 in B and T cells successfully identified a phenomenological model that could capture the synergy between the two cytokines.^25^ A cell population model has explained how T_reg_ IL-2 consumption suppresses effector T cell activation.^26^ However, any model needs to incorporate the key regulatory features of a pathway to accurately predict cell response. With structural information that clarified the mechanism of cytokine binding, for example, a model of IL-4, IL-7, and IL-21 binding revealed pathway cross-talk due to the relative γ_c_ receptor affinities.^27^ Nevertheless, these models have not accounted for endosomal trafficking nor been constructed to model multiple immune cell types. IL-2 induces rapid endocytosis-mediated IL-2Rα and IL-2Rβ downregulation,^14,28^ and trafficking is known to be a potent regulatory mechanism for all members of the γ_c_ family.^29^ Indeed, recent IL-15 engineering observed that attenuated cytokine potency can lead to *greater* therapeutic effect via reduced receptor-mediated clearance.^18^ Non-intuitive properties such as this can be better understood and optimized through models incorporating trafficking.

In this paper, we assemble a predictive model and tools to visualize γ_c_ cytokine family regulation. We first built a family-wide mathematical model that incorporates both binding and trafficking kinetics. This more comprehensive model allows us to investigate emergent behavior, such as competition between cytokines. This cytokine family is inherently high dimensional—with multiple ligands, cognate receptors, and cells with distinct expression. Therefore, we use tensor factorization to visualize the family-wide regulation. This map helps to identify how native or engineered ligands are targeted to specific immune cell populations based on their receptor expression levels. The methods used here can similarly be used in experimental and computational efforts of decoding other complex signaling pathways such as Wnt, Hedgehog, Notch, and BMP/TGFβ.^30–33^

## Results

### Trafficking is necessary to capture IL-2 and IL-15 dose response and the effect of IL-2Rα expression

To model how individual binding events give rise to cell response, we built a differential equation model representing the relevant binding and regulatory mechanisms within the γ_c_ receptor cytokine family (Fig. 1A). Binding interactions were modeled based on their known structural components, and led to the formation of receptor complexes capable of JAK/STAT signaling.^1^ Endocytic trafficking of cell surface receptors is a critical mechanism of regulatory feedback.^34–37^ Therefore, we extended earlier modeling efforts by including the trafficking of receptors and their complexes.^14,26^ We assumed that species trafficked into an endosomal compartment while continuing to produce JAK/STAT signaling and participate in binding events.

**Figure 1:**
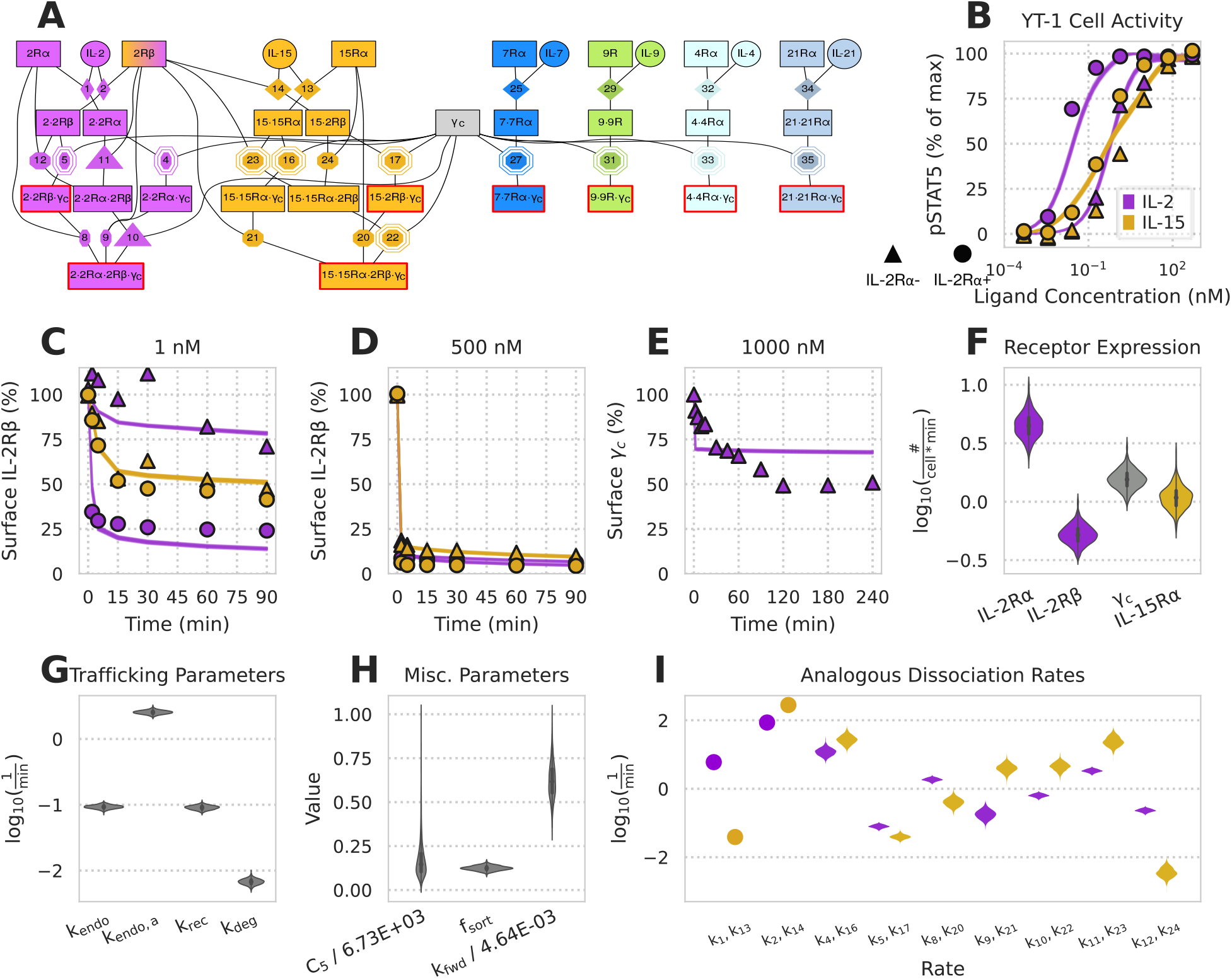
Incorporating trafficking leads to an accurate model of IL-2 & IL-15 response. A) Schematic of all receptor (boxes)-ligand (circles) complexes and binding events. Active (pSTAT signaling; containing two non-α receptors) complexes are outlined in red. Rate constants obtained from literature, detailed balance, or fitting are denoted by diamonds, octagons, or octagons with a double outline, respectively. Rate constants that were experimentally measured relative to other rates are denoted by triangles. B) Model prediction vs. experimental results for maximal pSTAT5 activation in YT-1 cells under various concentrations of ligand stimulation for 500 min. C-E) Model prediction vs. experimental results for the percent of initial IL-2Rβ (C, D) and γ_c_ (E) on the cell surface for various ligand stimulation concentrations and cell types. The 25-75% and 10-90% confidence intervals of model predictions are shaded dark and light respectively. Due to low prediction variability, only the 25-75% interval is visible. F-H) Posterior distributions after data fitting. C5 has units of # × cell^−1^, k_fwd_ has units of cell × #^−1^ × min^−1^, and f_sort_ is unitless. I) Posterior distributions for the analogous reaction rates of IL-2 and IL-15. Rates constants measured in literature are represented by dots.

Rate parameters for IL-2 and IL-15 binding events were parameterized by previous experimental measurements, detailed balance, or estimated by model fitting to existing experimental measurements (Fig. 1B–E). Fitting was performed to measurements of STAT5 phosphorylation and surface IL-2Rβ/γ_c_, upon either IL-2 or IL-15 stimulation, in either wild-type YT-1 human NK cells or YT-1 cells selected for expression of IL-2Rα. The posterior parameter distributions from these fits (Fig. 1F–I) were plugged back into our model and showed quantitative agreement with the data, including differential sensitivity with IL-2Rα expression (Fig. 1B–F).^14,38^ To evaluate the effect of including trafficking, we fit a version of the model without trafficking to the pSTAT5 measurements, using the same cell population as before; the model failed to fully capture differences with IL-2Rα expression even when using this limited fitting data (Fig. S1). Within the posterior distribution of parameter fits, IL-2·IL-2Rα complexes had a higher affinity for IL-2Rβ and γ_c_ than their IL-15oIL-15Rα counterparts in the trafficking model (k_4_ < k_16_ & k_11_ < k_23_), consistent with prior work (Fig. 1I).^39^ However, the opposite was inferred for IL-2Rβ (k_4_ > k_16_) and the affinities were equal for γ_c_ (k_11_ = k_23_) in the no-trafficking model (Fig. S1B). Depletion of surface IL-2Rβ and γ_c_ occurs through rapid endocytosis of active complexes and indeed, depletion occurred faster at higher cytokine doses (Fig. 1C-E). Correspondingly, active complex internalization (k_endo,a_) was inferred to be ~10x greater than that for inactive species (k_endo_) (Fig. 1G). These data indicated that accounting for trafficking is essential for modeling IL-2 and IL-15 signaling response.

Since IL-2 and IL-15 drive the formation of analogous active complexes, with IL-2Rβ, γ_c_, and a signaling-deficient high-affinity receptor (IL-2Rα/IL-15Rα), comparing their inferred binding rates gave insight into how IL-2 and IL-15 differ from one another (Fig.1I). The two ligands had nearly the same direct binding affinity to IL-2Rβ; however, IL-15 had a higher affinity than IL-2 for its α-chain. Consequently, IL-15’s complexes were inferred to more readily dimerize with a free α-chain than IL-2’s complexes (k_8_ > k_20_, k_12_ > k_24_). Similarly, IL-15 complexes had a slightly higher affinity for capturing IL-2Rβ/γ_c_ than their IL-2 counterparts (k_9_ < k_21_, k_10_ < k_22_, k_11_ < k_23_). The affinities of γ_c_ binding to ligand·IL-2Rβ and ligand·α-chain complexes were comparable between IL-2 and IL-15 (k_4_ =k_16_, k_5_ =k_17_). The data is also consistent with the literature in that both ligands have a higher affinity for IL-2Rβ when they are bound to their α-chain (k_2_, k_14_ > k_11_, k_23_).^39^ In total, a model of IL-2 and IL-15 incorporating trafficking is consistent with known biophysical and cell response measurements.

### Family model correctly captures IL-4/IL-7 dose responses and crossinhibition

To further test our model incorporating trafficking, we evaluated its performance in a series of experiments involving IL-4 and IL-7. IL-2 and IL-15 involve the same signaling-competent receptors and so the signaling activity of each cytokine cannot be distinguished. IL-4 and IL-7 activity, in contrast, can be distinguished when both cytokines are co-administered to cells by measuring STAT6 and STAT5 phosphorylation, respectively.^2^ Using this phenomenon we explored cross-inhibition data wherein IL-4 and IL-7doses were administered to human PBMC-derived T cells (CD4^+^TCR^+^CCR7^high^) both individually and together.^27^

Using surface abundance measurements ofIL-4Rα, IL-7Rα, and γ_c_, we applied a steadystate assumption in the absence of ligand to solve for each receptor expression rate.^27^ Our model fits both single and dual cytokine dose-response data sets with high accuracy (Fig. 2B–C). The fitting process identifiably constrained reaction rates, trafficking parameters, and pSTAT scaling constants (Fig. 2F–I). While surface abundance was constrained, the receptor expression rates still formed distributions dependent on trafficking parameters (Fig. 2G–I).

**Figure 2:**
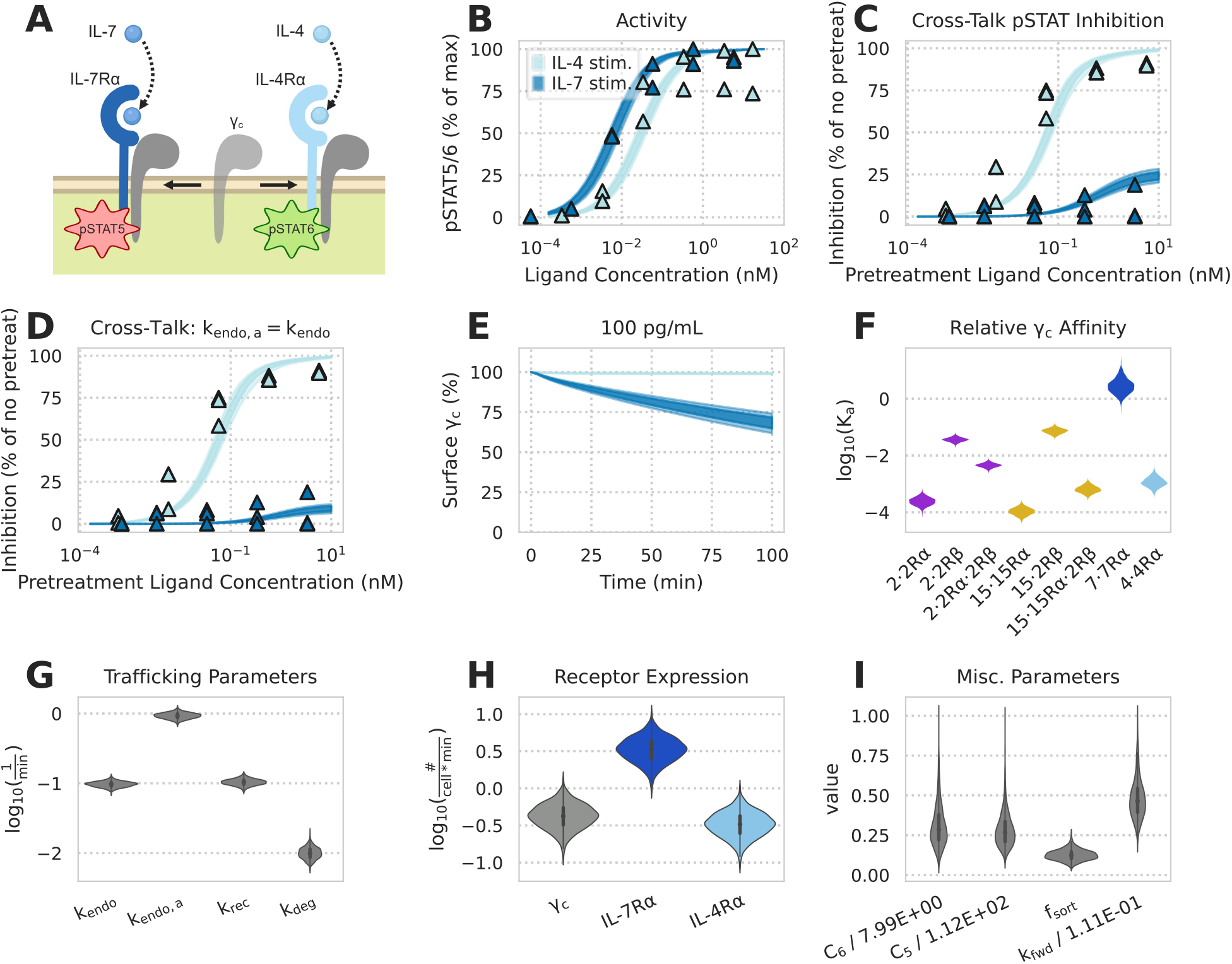
A reaction model captures cytokine-cytokine interactions. A) Schematic of IL-4 and IL-7 receptor complexes competing for γ_c_ and generating distinct pSTAT signals. B-C) Fitting model to experimental data. Experimental measurements are denoted by triangles. Shaded areas represent the 25-75% and 10-90% confidence intervals of model predictions. pSTAT5 and pSTAT6 were measured for IL-7 and IL-4 experiments, respectively. B) Single-cytokine pSTAT dose-response measurements for 10 min of exposure to IL-4 and IL-7. C) Percent inhibition of the second cytokine’s pSTAT response in a dual-cytokine dose-response experiment. Human PBMC-derived T cells (CD4^+^TCR^+^CCR7^high^) were pretreated with various concentrations of one cytokine for 10 min before being stimulated with a fixed concentration (50 pg/mL IL-7 or 100 pg/mL IL-4) of the other cytokine for an additional 10 min. D) Model predictions for percent inhibition of the second cytokine’s pSTAT response in a dual-cytokine dose-response experiment with the assumption that active species are endocytosed at the same rate as inactive species (k_endo,a_ = k_endo_). E) Model predictions for percent of γ_c_ on the cell surface when exposed to 100 pg/mL of either IL-7 or IL-4 for 100 min. F) Violin plot of K_a_ values obtained via posterior distributions of k_fwd_ / k_rev_ for krev parameters corresponding to different complexes competing for the common γ_c_ (Fig. 1A). G–I) Posterior distributions from fitting to data. Scaling constants C_5_ and C_6_ have units of # × cell^−1^, k_fwd_ has units of cell × #^−1^ × min^−1^, and f_sort_ is unitless

The experimental data and model fits showed that IL-7inhibited IL-4activity more than vice versa (Fig. 2C).^27^ Consistent with the experimentally-derived mechanism,^27^ this inhibitory behavior was explained by the competition of ligandoα-chain complexes for the common γ_c_. The inferred Kd of this dimerization process for IL-7 (k_27_) was smaller than the K_d_ for IL-4 (k_33_), indicating that there was tighter dimerization of IL-7oIL-7Rα to γ_c_ than there was dimerization of IL-4·IL-4Rα to γ_c_ (Fig. 2F). The competition for γ_c_ was determined to play a larger role in signaling inhibition than receptor internalization since our model predicted that the same inhibitory relationships hold when active complexes internalize at the same rate as other species (Fig. 2D). Internalization was additionally dismissed because the majority of γ_c_ remained on the cell surface after ligand stimulation in both model simulation and experimental measurement (Fig. 2E).^27^

### Tensor Factorization Maps the Gamma Chain Family Response Space

Even with an accurate model, exploring how dynamic responses vary across responding cell types and ligand treatments remains challenging. Restricting ones’ view to a single time point, cell type, or ligand concentration provides only a slice of the picture. Therefore, we sought to apply factorization as a means to globally visualize ligand response.

As response to ligand is mostly defined by receptor expression, we quantitatively profiled the abundance of each IL-2, IL-15, and IL-7 receptor across ten PBMC subpopulations (Fig. 3A). PBMCs were stained using receptor-specific fluorescent antibodies and analyzed by flow cytometry; their subpopulations were separated using canonical markers (Fig. S3, tbl. S1). These data recapitulated known variation in these receptors, including high IL-7Rα or IL-2Rα expression in helper and regulatory T cells, respectively.^1,40^ As mentioned above, IL-7 is uniquely able to cross-inhibit other γ_c_ cytokines, and excess IL-7Rα likely helps to ensure this occurs (Fig. 2C).^27^ Principal component analysis (PCA) helped visualize variation in this receptor abundance data (Fig. 3B-C). Principal component 1 most prominently separated the NK cells from all others due to their distinct receptor expression, with high levels of IL-2Rβ and relatively lower levels ofγ_c_. Principal component 2 then separated effector and regulatory T cell populations, based on their high IL-7Rα or IL-2Rα abundance, respectively. However, PCA also helped to identify slightly higher γ_c_ levels in Tregs, and the slightly more Treg-like profile of memory CD8+ cells.

**Figure 3:**
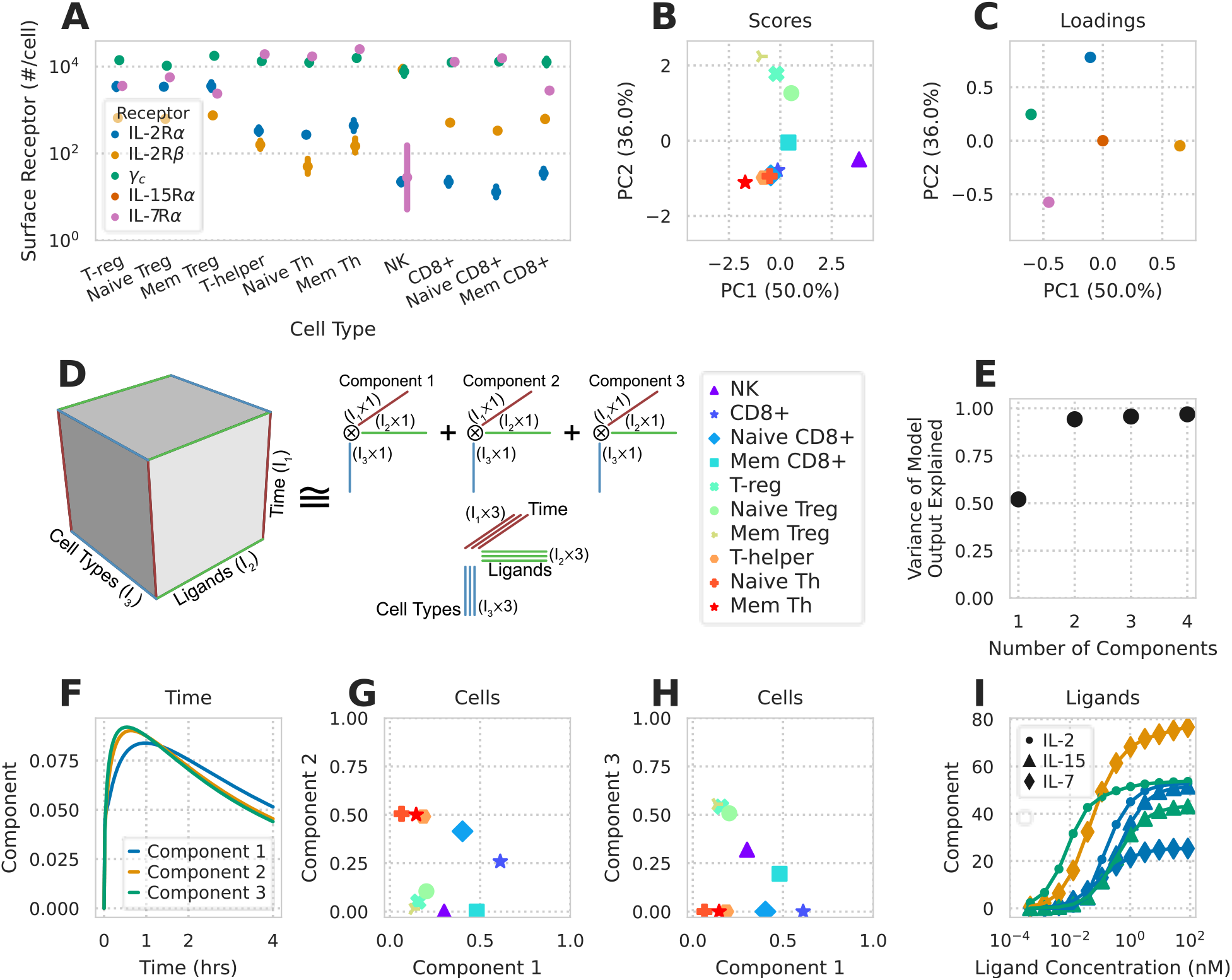
Tensor factorization to map model-predicted cytokine responses. A) Measured receptor abundance for ten PBMC-derived subpopulations. Points and error bars show geometric mean and standard deviation respectively (N = 4). Error bars for some points are too small to display. B-C) PCA scores (B) and loadings (C) of receptor abundance. Axis label percentages indicate percent variance explained. D) Schematic representation of CP decomposition. Model predictions are arranged in a cube depending upon the time, ligand treatment, and cell type being modeled. CP decomposition then helps to visualize this space. E) Percent variance reconstructed (R2X) versus the number of components used in non-negative CP decomposition. F-I) Component values versus time (F), cell type (G-H), or ligand stimulation (I). The variation explained by each component is the product of the component’s time, ligand, and cell type factorization. Ligand components with only negligible values (< 5% max) are not shown.

To build a tensor of model predictions, we assembled simulation predictions across cell types, ligand conditions, and time. This three-dimensional (time, cell type, ligand) tensor was then decomposed with non-negative canonical polyadic (CP) decomposition (Fig. 3D). We selected three components during decomposition as this number captured 95% of the variance in our original data tensor (Fig. 3E). To show the relationships among the tensor’s three dimensions, the component plots of each dimension were plotted alongside each other.

CP decomposition can be interpreted by matching a single component’s effects across factor plots for each dimension. For example, component 2 is greatest at roughly 50 mins, for helper and CD8+T cells, and almost exclusively with IL-7 stimulation (Fig. 3F– I). This indicates that this variation in the data occurs with IL-7 stimulation, leads to a response in helper and CD8+ T cells, and peaks at 50 mins. In this way, different contributory factors in cell response are separated.

All components showed similar variation with time, peaking quickly and then decreasing after roughly 50 mins (Fig. 3F). This can be understood through two phases, in which receptor activation occurs, and then trafficking-mediated downregulation of the receptors (Fig. 1). Comparing the cells and ligands decomposition plots showed expected effects. IL-7 response was separated as component 2, showed a dose-dependent increase, and correlated with IL-7Rα expression levels (Fig. 3A/G/I). Interestingly, IL-2/-15 response separated by concentration, rather than ligand. Low concentrations of IL-2 were represented by component 3, and preferentially activated regulatory over effector Tcells (Fig. 3H/I). High concentrations of IL-2/-15 were represented by component 1 and similarly activated effector and regulatory T cells (Fig. 3G/I). This known dichotomy occurs through higher IL-2Rα expression in T_reg_s (Fig. 3A). Importantly, while PCA can help to distinguish cells based on distinct receptor expression profiles, cells separated differently based on their predicted ligand stimulation response (Fig. 3B/G/H). This demonstrates the unique benefit of tensor-and model-based factorization to distinguish cells based upon their predicted response profiles.

Other tensor decomposition methods exist and can also be applied to visualize model-predicted response. For example, non-negative Tucker decomposition relaxes CP decomposition by employing a core tensor enabling interaction terms between components (Fig. S4).^41^ However, this flexibility comes at the cost of interpretability, as visualizing the core tensor’s effect is challenging. In total, factorization methods provide an effective means of visualizing the high-dimensional regulation of complex receptor families, including the influence of time, ligand stimulation, and receptor expression.

### Accurately Predicted Response Across a Panel of PBMC-Derived Cell Types

We evaluated whether our model accurately predicts differences in the cell type-specificity of ligand treatment by comparing its predictions for IL-2/-15 responses across a panel of 10 PBMC-derived cell populations. We both measured and used our model to predict PBMC response to cytokine stimulation at 12 concentrations (0.5 pM–84 nM) and 4 time points (30 minutes, 1, 2, and 4 hours). Individual cell types displayed reproducible responses to IL-2/-15 treatment (Fig. 4A). Overall, our model predictions of ligand pSTAT5 response closely matched experimental measurement (Figs. 4, S5). The differences between cell types largely matched known differences in cytokine response. For example, T_reg_s were markedly sensitive to IL-2 (Fig. 4B/F), but not IL-15 (Fig. 4B/I), at low concentrations of the cytokine.^23,24^ Small amounts of of IL-2Rα in helper T cells (Fig. 3A) partially sensitizes them to IL-2 (Fig. 4B; Fig. S5H). Our model accurately captured these differences in sensitivity and response across all the cell populations (Fig. 4C).

**Figure 4:**
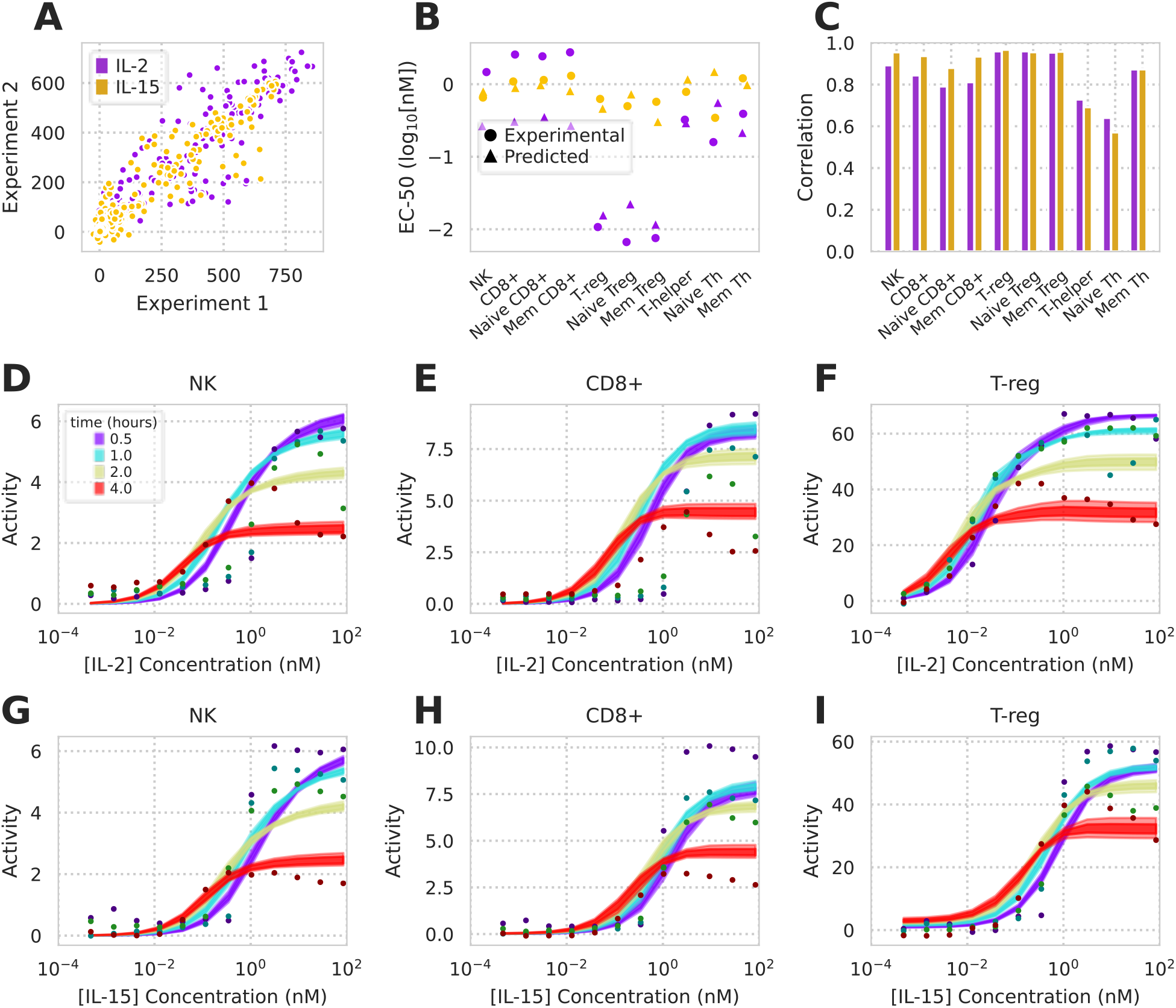
Model accurately predicts cell type-specific response across a panel of PBMC-derived cell types. A) Comparison of two replicates measuring pSTAT5 response to a dose-response of IL-2/-15, time course, and panel of PBMC-derived cell types. B) Both experimentally-derived and model-predicted EC_50_s of dose response across IL-2/-15 and all 10 cell types. EC_50_s are shown for 1 hr time point. C) Pearson correlation coefficients between model prediction and experimental measurements for all 10 cell populations (full data shown in Fig. S5). D–I) pSTAT5 response to IL-2 (D-F) or IL-15 (G-I) dose responses in NK, CD8+, and T_reg_ cells.

While the model accurately predicted experimentally-measured responses overall, and specifically the sensitivities of the dose-response profiles, we noticed some discrepancy specifically at high ligand concentrations and longer times in specific cell populations (Fig. 4; Fig. S5). For example, while CD8+ cells almost exactly match model predictions at 1 hr, by 4 hrs we experimentally observed a biphasic response with respect to IL-2 concentration, and a plateau with IL-15 that decreased over time. This decrease in signaling was most pronounced with the CD8+ cells, but could be observed to lesser extents in some other cell populations such as NK cells (Fig. S5). We hypothesize two possible explanations for this discrepancy: First, CD8+ populations are known to proteolytically shed IL-2Rα in an activity-responsive manner.^42^ Second, our model only uses a very simple sigmoidal relationship between active receptor and pSTAT5 signal. Other components of the JAK-STAT pathway surely influence its dynamic response.^43^ However, overall the model presented here remains useful for exploring the determinants of cell type-specific response, which originate at the receptor expression profile on the cell surface.

### Tensor Factorization of Experimental Measurements Distinguishes Cell Type-Specific Responses

Given that tensor factorization helped to visualize model predictions of IL-2, −7, and −15 response, we wished to evaluate whether it could similarly help to visualize experimental measurements. We structured our experimental pSTAT5 measurements in an identical format to the model simulation tensor Fig. 3. Factoring into two components explained roughly 90% of the variance in the original data (Fig. 5A), which we can then interpret using each of the factor plots (Fig. 5B–D).

**Figure 5:**
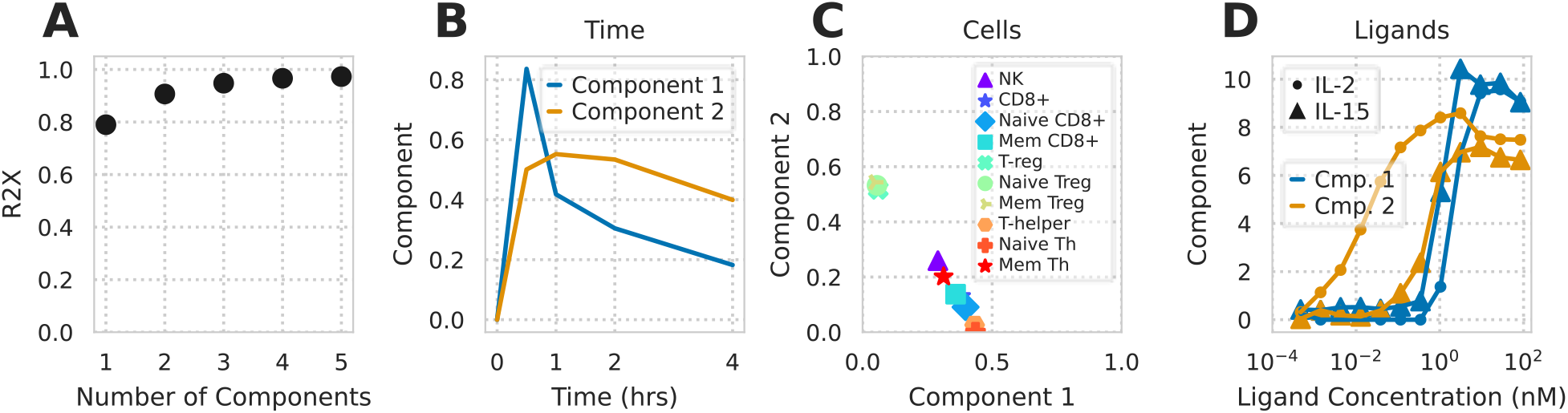
Non-negative CP decomposition applied to experimental pSTAT5 measurements. A) R2X of non-negative CP decomposition versus number of components used. B–D) Decomposition plot with respect to time (B), cell type (C), or ligand treatment (D).

Interestingly, these factors are distinguished by their concentration dependence more so than being tied to a specific ligand (Fig. 5D). Component 2 increases with low concentrations of IL-2, while component 1 only increases at high concentrations of either ligand. As expected, effector and regulatory T cells are most strongly associated with components 1 and 2, respectively, matching their known dose-response profiles (Fig. 4). However, component 2 is also distinct from 1 in its sustained activation (Fig. 5B; Fig. S5). This can be expected from rapid endocytosis-mediated downregulation of IL-2Rβ at high IL-2/-15 concentrations (Fig. 1). Thus, tensor factorization helps to separate these differences in dose-and cell type-specific responses. Furthermore, there was clear correspondence between the model and experimental factorization. For example, the low-dose IL-2-specific component in the model and experiment factorization correlated strongly in their cell type weighting (cosine similarity of 0.96; Fig. 3H; Fig. 5C).

### Model Accurately Captures Cell Type-Specific Response to IL-2 Muteins

Using the model, we sought to identify strategies for selectively targeting T_reg_s. In order to quantify the effectiveness of selectively activating T_reg_s, we defined a specificity metric as the normalized pSTAT5 response of T_reg_s divided by the pSTAT5 response ofT-helper or NK cells. As expected, both model prediction and experimental values of this specificity increased with lower concentrations of IL-2 and had a lesser concentration dependent relationship with IL-15 (Fig. 6A/B). With this quantity, we then examined the sensitivity of the specificity metric with respect to both surface and endosomal binding. Decreasing IL-2Rα unbinding (k5rev), particularly in the endosome, provided the largest and most consistent benefit to specificity (Fig. 6C). Changes in endosomal binding rates have been shown to have important effects on protein therapy’s half-life and potency.44 To the extent this binding can be separately manipulated, the model indicates it might help to improve specificity as well. Moreover, the model predicts that ligands with reduced IL-2Rα affinity had a decreased ability to specifically activate T_reg_s with respect to NK and T-Helper cells regardless of their IL-2Rβ/γ_c_ affinity (Fig. 6D). Therefore, while reducing IL-2Rβ/γ_c_ affinity can help modulate the potency of these cytokines, maintaining IL-2Rα affinity may be especially critical. In total, these results demonstrate this model’s ability to predict immune cell response to wild-type or engineered cytokines, particularly for engineering cell-specific responses.

**Figure 6:**
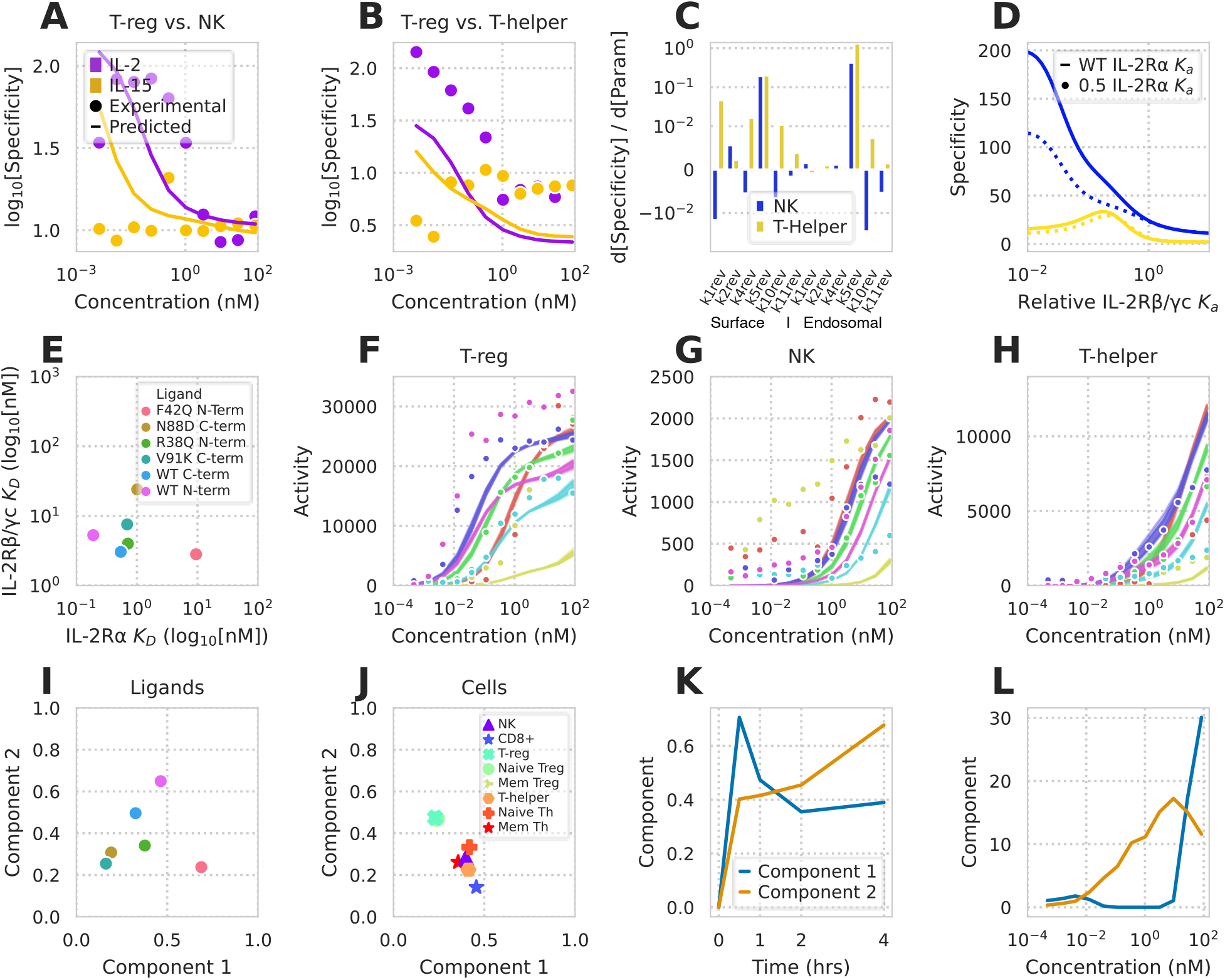
Model and tensor factorization predicts and decodes cell type-specific responses to IL-2 muteins. A-B) Predicted and measured T_reg_ activation specificity compared to NK (A) and T helper (B) cells. C) Partial derivatives of T_reg_ activation specificity compared to NK and T helper cells with respect to each surface and endosomal reverse binding rate constant. D) T_reg_ activation specificity with respect to NK and T helper cells as a function of IL-2Rβ/γc binding affinity for ligands with wild type and reduced IL-2Rα affinity. Specificity values are shown for cells exposed to a cytokine concentration of 38 pM. E) IL-2Rα and IL-2Rβ/γc dissociation constants for our panel of IL-2 muteins. F-H) Predicted versus experimental immune cell responses to IL-2 muteins for T_reg_s (F), NK cells (G), and T-helpers (H). Dots represent experimental measurements and shaded regions represent 10-90% confidence interval for model predictions. Mutein stimulant denoted by color. I-L) Tensor factorization of experimentally measured cellular activation values for IL-2 muteins. Component values versus ligand (I), cell type (J), time (K), and cytokine concentration (L).

To evaluate the potential of the model for cytokine engineering, we measured PBMC response to several Fc-bound IL-2 monomers. Several wild-type and mutant forms of IL-2 were produced as fusions with a monomeric human antibody Fc domain. Targeted mutations were introduced to IL-2 domains known to be instrumental to either IL-2Rα or IL-2Rβ/γ_c_ binding. Cytokines are often Fc-conjugated to increase the drug’s *in vivo* half-life, and can be conjugated in a variety of orientations. We quantified the effect of our engineered mutations and Fc conjugation on IL-2Rα and IL-2Rβ/γ_c_ binding kinetics using bio-layer inferometry (Fig. S6). Surprisingly, we found that Fc-conjugation to the N-terminus selectively lowered IL-2Rβ/γ_c_ affinity, while conjugation to the C-terminus selectively lowered IL-2Rα affinity (tbl. SD1, Fig. 6E). Therefore, Fc conjugation can have either complementary or counterproductive effects on mutation-mediated changes in receptor affinity, and affinity must be assessed in the clinical format.

Using these altered affinities, we were able to accurately predict cell type-specific activity response to our modified ligands (Fig. S7, Fig. 6F-H). Ligands with decreased IL-2Rα or IL-2Rβ/γ_c_ affinity had decreased T_reg_ or T-Helper activity response, respectively, as expected. As before, visualizing the effect of altered binding kinetics on cellular response is complicated by the contribution of cell type, concentration, and time (Fig. 3E-I). In order to visualize our results, we performed tensor factorization using the experimentally-determined pSTAT5 response of PBMCs exposed to both wild-type and modified IL-2 ligands (Fig. 6I-L). Two components explained 80% of the variance in the new combined data tensor. Among the ligands, wild-type N-terminally conjugated IL-2 was the most potent inducer of T_reg_ response as shown by its strong component 2 weighting (Fig. 6I/J). The difference in signaling with Fc conjugation orientation is likely due to these conjugation types’ opposing effects on the cytokine’s IL-2Rα affinity.

## Discussion

Here, we built a mass-action kinetic binding model for the common γ_c_ receptor family, and used factorization methods to explore its cell type-dependent behavior. This approach provided insights into its high-dimensional regulation. Our binding-reaction model combined the structure of ligand interaction with endosomal trafficking, both of which were critical for accurately modeling response (Fig. 1 & Fig. S1). After fitting our model to previously published cytokine response data, we were able to predict IL-2 and −15 response across a wide panel of PBMC-derived cell types (Fig. 4). Mass-action models can help to explain counter-intuitive features of ligand response and identify specific strategies for optimizing therapeutically-desired properties.^45,46^ In the case of the γ_c_ receptor cytokines, a therapeutic goal has been to specifically modulate subpopulations of cells based on their unique receptor expression profiles.^12,22–24^ To visualize these possibilities, we employed tensor factorization to map the signaling response space. This map provided a clearer picture of differential responsiveness between ligands, with selective and increased activation for certain cells and ligands (Fig. 5 & fig. 6). For example, we could clearly identify the selectivity of T helper cells for IL-7, and low concentrations of IL-2 for T_reg_s (Fig. 3).

The model described here serves as an effective tool for cell type-selective rational cytokine design. In addition to the natural ligands, many cytokine muteins have been designed with altered binding affinities to specific receptors.^20,21^ Our model serves as a computational tool for comparing these muteins as immunotherapeutic drugs that selectively activate certain cell populations. For example, our model helped to identify that high IL-2Rα affinity is essential to preserve T_reg_ specificity, regardless of the affinity toward IL-2Rβ/γ_c_ (Fig. 6). Fc conjugation orientation can significantly influence receptor affinity (including reducing IL-2Rα affinity), and so this step of drug design needs to be incorporated into ligand optimization (Fig. 6E). Incorporating trafficking with the binding events of the cytokines allowed us to distinguish surface and endosomal binding, which is an unexplored axis for further engineering cell-specific responses. Indeed, endosomal IL-2Rα affinity is predicted to be *more* critical to T_reg_ specificity than binding on the surface, which agrees with the distinct temporal profiles of ligand response between cell types on the time-scale of trafficking (Fig. 6C & K).

Models incorporating the full panel of responding cell populations will enable further refinement of these engineered ligands.^47^ Both IL-2 and IL-15 have extremely short half-lives *in vivo*, in part due to endocytosis mediated clearance.^17,18^ Including endocytic trafficking of ligand will enable future work modeling ligand clearance *in vitro* and *in vivo*. Changes in receptor binding may therefore be selected based on both optimized selectivity and pharmacokinetic properties. While cell types were defined here by their average receptor expression, cell-to-cell variability within these populations leads to variation in stimuli response.^15^ Incorporating single cell variation will provide a more complete picture of population response, and may help to further refine cell type selectivity.

Receptor families with many receptors and ligands are often made up of a dense web of connections, making the role of individual components non-intuitive.^30,33^ Interconnected, cross-reactive components may have evolved as a tradeoff between transmitting ligand-mediated information and expanding the repertoire of cell-surface proteins.^48^ The methods detailed in this paper can be applied to many signaling systems characterized by pleiotropy and high-dimensionality. The combination of dynamical, mechanistic models and statistical exploration methods is particularly powerful to provide actionable directions for how to optimize therapeutic response. Detailed biophysical and structural characterization, animal disease models, and evidence from human genetic studies make this engineering possible for therapeutically targeting other other complex signaling pathways including FcγR, Wnt, Hedgehog, Notch, and BMP/TGFβ.^30–33,49^

## Methods

All analysis was implemented in Python, and can be found at https://github.com/meyer-lab/gc-cytokines, release 1.0.

### Model

#### Base model

Cytokine (IL-2, −4, −7, −9, −15, & −21) binding to receptors was modeled using ordinary differential equations (ODEs). IL-2 and −15 each had two private receptors, one being a signaling-deficient α-chain (IL-2Rα & −15Rα) and the other being signaling-competent IL-2Rβ. The other four cytokines each had one signaling-competent private receptor (IL-7Rα, −9R, −4Rα, & −21Rα). JAK-STAT signaling is initiated when JAK-binding motifs are brought together. JAK binding sites are found on the intracellular regions of the γ_c_, IL-2Rβ, IL-4Rα, IL-7Rα, IL-9R, and IL-21Rα receptors; therefore all complexes which contained two signaling-competent receptors were deemed to be active species. Ligands were assumed to first bind a private receptor and then can dimerize with other private receptors or γ_c_ thereafter. Direct binding of ligand to γ_c_ was not included due to its very weak or absent binding.^50^

In addition to binding interactions, our model incorporated receptor-ligand trafficking. Receptor synthesis was assumed to occur at a constant rate. The endocytosis rate was defined separately for active (k_endo,a_) and inactive (k_endo_) receptors. f_sort_ fraction of species in the endosome were ultimately trafficked to the lysosome, and active species in the endosome had a sorting fraction of 1.0. All endosomal species not sent to lysosomes were recycled back to the cell surface. The lysosomal degradation and recycling rate constants were defined as k_deg_ and k_rec_, respectively. We assumed no autocrine ligand was produced by the cells. We assumed an endosomal volume of 10 fL and endosomal surface area half that of the plasma membrane.^46^ All binding events were assumed to occur with 5-fold greater disassociation rate in the endosome due to its acidic pH.^34^

Free receptors and complexes were measured in units of number per cell and soluble ligands were measured in units of concentration (nM). Due to these unit choices for our species, the rate constants for ligand binding to a free receptors had units of nM^−1^ min^−1^, rate constants for the forward dimerization of free receptor to complex had units of cell min^−1^ number^−1^. Dissociation rates had units of min^−1^. All ligand-receptor binding processes had an assumed forward rate (k_bnd_) of 10^7^ M^−1^ sec^−1^. All forward dimerization reaction rates were assumed to be identical, represented by k_fwd_. Reverse reaction rates were unique. Experimentally-derived affinities of 1.0,^27^ 59,^51^ 0.1,^52^ and 0.07 nM^27^ were used for IL-4, −7, −9, and −21 binding to their cognate private receptors, respectively. IL-2 and −15 were assumed to have affinities of 10 nM and 0.065 nM for their respective α-chains,^53–55^ and affinities of 144 nM and 438 nM for their respective β-chains.^53^ Rates k_5_, k_10_, and k_11_ were set to their experimentally-determined dissas-sociation constants of 1.5, 12, and 63 nM.^53^

Initial values were calculated by assuming steady-state in the absence of ligand. Differential equation solving was performed using the SUNDIALS solvers in C++, with a Python interface for all other code.^56^ Model sensitivities were calculated using the adjoint solution.^57^ Calculating the adjoint requires the partial derivatives of the differential equations both with respect to the species and unknown parameters. Constructing these can be tedious and error-prone. Therefore, we calculated these algorithmically using forward-pass autodifferentiation implemented in Adept-2.^58^ A model and sensitivities tolerance of 10^−9^ and 10^−3^, respectively, were used throughout. We used unit tests for conservation of mass, equilibrium, and detailed balance to help ensure model correctness.

#### Model fitting

We used Markov chain Monte Carlo to fit the unknown parameters in our model using previously published cytokine response data.^14,27^ Experimental measurements include pSTAT activity under stimulation with varying concentrations of IL-2, −15, −4, and −7 as well as time-course measurements of surface IL-2Rβ upon IL-2 and −15 stimulation. YT-1 human NK cells were used for all data-sets involving IL-2 and IL-15. Human PBMC-derived CD4+TCR+CCR7^high^ cells were used for all IL-4 and −7 response data. All YT-1 cell experiments were performed both with the wild-type cell line, lacking IL-2Rα, and cells sorted for expression of the receptor. Data from Ring *et al* and Gonnord *et al* can be found in Figure 5 and Figure S3 of each paper, respectively.^14,27^ Measurements of receptor counts at steady state in Gonnord *et al* were used to solve for IL-7Rα, IL-4Rα, and γ_c_ expression rates in human PBMCs.

Fitting was performed with the Python package PyMC3. All unknown rate parameters were assumed to have a lognormal distribution with a standard deviation of 0.1; the only exception to these distributions was f_sort_ which was assumed to have a beta distribution with shape parameters of α=20 and β=40. Executing this fitting process yielded likelihood distributions of each unknown parameter and sum of squared error between model prediction and experimental data at each point of experimental data. The Geweke criterion metric was used to verify fitting convergence for all versions of the model (Fig. S2).^59^

#### Tensor Generation and Factorization

To perform tensor factorization we generated a three-(timepoints × cell types × ligand) or four-dimensional (timepoints × cell types × concentration × mutein) data tensor of predicted or measured ligand-induced signaling. Before decomposition, the tensor was variance scaled across each cell population. Tensor decomposition was performed using the Python package TensorLy.^60^ Except where indicated otherwise, tensor decomposition was performed using non-negative canonical polyadic decomposition. Where indicated, non-negative Tucker decomposition was used.

### Experimental Methods

#### Receptor abundance quantitation

Cryopreserved PBMCs (ATCC, PCS-800-011, lot#81115172) were thawed to room temperature and slowly diluted with 9 mL pre-warmed RPMI-1640 medium (Gibco, 11875-093) supplemented with 10% fetal bovine serum (FBS, Seradigm, 1500-500, lot#322B15). Media was removed, and cells washed once more with 10 mL warm RPMI-1640 + 10% FBS. Cells were brought to 1.5×10^6^ cells/mL, distributed at 250,000 cells per well in a 96-well V-bottom plate, and allowed to recover 2 hrs at 37°C in an incubator at 5% CO2. Cells were then washed twice with PBS + 0.1% BSA (PBSA, Gibco, 15260-037, Lot#2000843) and suspended in 50 μL PBSA + 10% FBS for 10 min on ice to reduce background binding to IgG.

Antibodies were diluted in PBSA + 10% FBS and cells were stained for 1 hr at 4°C in darkness with a gating panel (Panel 1, Panel 2, Panel 3, or Panel 4) and one antireceptor antibody, or an equal concentration of matched isotype/fluorochrome control antibody. Stain for CD25 was included in Panel 1 when CD122, CD132, CD127, or CD215 was being measured (CD25 is used to separate T_reg_s from other CD4+ T cells).

Compensation beads (Simply Cellular Compensation Standard, Bangs Labs, 550, lot#12970) and quantitation standards (Quantum Simply Cellular anti-Mouse IgG or anti-Rat IgG, Bangs Labs, 815, Lot#13895, 817, Lot#13294) were prepared for compensation and standard curve. One well was prepared for each fluorophore with 2 μL antibody in 50 μL PBSA and the corresponding beads. Bead standards were incubated for 1 hr at room temperature in the dark.

Both beads and cells were washed twice with PBSA. Cells were suspended in 120 μL per well PBSA, and beads to 50 μL, and analyzed using an IntelliCyt iQue Screener PLUS with VBR configuration (Sartorius) with a sip time of 35 and 30 secs for cells and beads, respectively. Antibody number was calculated from fluorescence intensity by subtracting isotype control values from matched receptor stains and calibrated using the two lowest binding quantitation standards. T_reg_ cells could not be gated in the absence of CD25, so CD4+ T cells were used as the isotype control to measure CD25 in T_reg_ populations. Cells were gated as shown in Fig. S3. Measurements were performed using four independent staining procedures over two days. Separately, the analysis was performed with anti-receptor antibodies at 3x normal concentration to verify that receptor binding was saturated. Replicates were summarized by geometric mean.

#### pSTAT5 Measurement of IL-2 and -15 Signaling in PBMCs

Human PBMCs were thawed, distributed across a 96-well plate, and allowed to recover as described above. IL-2 (R&D Systems, 202-IL-010) or IL-15 (R&D Systems, 247-ILB-025) were diluted in RPMI-1640 without FBS and added to the indicated concentrations. To measure pSTAT5, media was removed, and cells fixed in 100 μL of 10% formalin (Fisher Scientific, SF100-4) for 15 mins at room temperature. Formalin was removed, cells were placed on ice, and cells were gently suspended in 50 μL of cold methanol (−30°C). Cells were stored overnight at −30°C. Cells were then washed twice with PBSA, split into two identical plates, and stained 1 hr at room temperature in darkness using antibody panels 4 and 5 with 50 μL per well. Cells were suspended in 100 μL PBSA per well, and beads to 50 μL, and analyzed on an IntelliCyt iQue Screener PLUS with VBR configuration (Sartorius) using a sip time of 35 seconds and beads 30 seconds. Compensation was performed as above. Populations were gated as shown in Fig. S3, and the median pSTAT5 level extracted for each population in each well.

#### Recombinant proteins

IL-2/Fc fusion proteins were expressed using the Expi293 expression system according to manufacturer instructions (Thermo Scientific). Proteins were as human IgG1 Fc fused at the N-or C-terminus to human IL-2 through a (G4S)4 linker. C-terminal fusions omitted the C-terminal lysine residue of human IgG1. The AviTag sequence GLNDIFEAQKIEWHE was included on whichever terminus did not contain IL-2. Fc mutations to prevent dimerization were introduced into the Fc sequence.^61^ Proteins were purified using MabSelect resin (GE Healthcare). Proteins were biotinylated using BirA enzyme (BPS Biosciences) according to manufacturer instructions, and extensively buffer-exchanged into phosphate buffered saline (PBS) using Amicon 10 kDa spin concentrators (EMD Millipore). The sequence of IL-2Rβ/γ Fc heterodimer was based on a reported active heterodimeric molecule (patent application US20150218260A1), with the addition of (G4S)2 linker between the Fc and each receptor ectodomain. The protein was expressed in the Expi293 system and purified on MabSelect resin as above. IL2-Rα ectodomain was produced with C-terminal 6xHis tag and purified on Nickel-NTA spin columns (Qiagen) according to manufacturer instructions.

#### Octet binding assays

Binding affinity was measured on an OctetRED384 (ForteBio). Briefly, biotinylated monomeric IL-2/Fc fusion proteins were uniformly loaded to Streptavidin biosensors (ForteBio) at roughly 10% of saturation point and equilibrated for 10 minutes in PBS + 0.1% bovine serum albumin (BSA). Association time was up to 40 minutes in IL-2Rβ/γ titrated in 2x steps from 400 nM to 6.25 nM, or IL-2Rα from 25 nM to 20 pM, followed by dissociation in PBS + 0.1% BSA. A zero-concentration control sensor was included in each measurement and used as a reference signal. Assays were performed in quadruplicate across two days. Binding to IL-2Rα did not fit to a simple binding model so equilibrium binding was used to determine the K_D_ within each assay. Binding to IL-2Rβ/γ fit a 1:1 binding model so on-rate (k_on_), off-rate (k_off_) and K_D_ were determined by fitting to the entire binding curve. Kinetic parameters and K_D_ were calculated for each assay by averaging all concentrations with detectable binding signal (typically 12.5 nM and above).

## Acknowledgements

This work was supported by NIH DP5-OD019815 to A.S.M. and by a research agreement with Visterra Inc.

## Author contributions statement

A.S.M. and S.M.C. conceived of the study. S.M.C. and C.P. performed the PBMC experiments and engineered the IL-2 fusion proteins. A.C.W., A.M.F., A.S.M, B.O.J., and Z.S.K. performed the computational analysis. All authors helped to design experiments and/or analyze the data.

## Supplement

### IL-2, IL-15, and IL-7 Receptor Quantitation

**Table S1:** Antibodies used to quantify receptors and cell types. *Panel 0:* Antibodies for IL-2, IL-15, and IL-7 receptor analysis; *Panel 1:* Antibodies to gate Naïve and Memory T-regulatory and T-helper cells; *Panel 2:* Antibodies to gate NK and CD56bright NK cells; *Panel 3:* Antibodies to gate Naïve and Memory Cytotoxic T cells; *Panel 4:* Antibodies to gate Naïve and Memory T-regulatory, T helper, and Cytotoxic cells, and NK cells for CD127 (IL-7) Quantitation; *Panel 5:* Antibodies to gate Memory and Naïve T-regulatory cells, Memory and Naïve T-helper cells; *Panel 6:* Antibodies to gate NK cells, CD56bright NK cells, and Cytotoxic T cells. ^⋆^CST: Cell Signaling Technology.

| Antibody (clone) | Dilution | Fluorophore | Vendor (CAT#) | Panel |
| --- | --- | --- | --- | --- |
| CD25 (M-A251) | 1:120 | Brilliant Violet 421 | BioLegend (356114) | 0 |
| CD122 (TU27) | 1:120 | PE/Cy7 | BioLegend (339014) | 0 |
| CD132 (TUGh4) | 1:120 | APC | BioLegend (3386) | 0 |
| CD215 1st mAb (JM7A4) | 1:120 | APC | BioLegend (330210) | 0 |
| CD215 2nd mAb (151303) | 3:100 | APC | R&D Systems (FAB1471A) | 0 |
| CD127 (A019D5) | 1:120 | Alexa Fluor 488 | BioLegend (351313) | 0 |
| Ms IgG1κ (MOPC-21) | 1:240 | Brilliant Violet 421 | BioLegend (400158) | 0 |
| Md IgG1κ (MOPC-21) | 1:240 | PE/Cy7 | BioLegend (400126) | 0 |
| Rat IgG2Bκ (RTK4530) | 1:60 | APC | BioLegend (400612) | 0 |
| Ms IgG2Bκ (MPC-11) | 1:120 | APC | BioLegend (400320) | 0 |
| Ms IgG2B (133303) | 3:100 | APC | R&D Systems (IC0041A) | 0 |
| Ms IgG1κ (MOPC-21) | 1:120 | Alexa Fluor 488 | BioLegend (400129) | 0 |
| CD3 (UCHT1) | 1:120 | Brilliant Violet 605 | BioLegend (300460) | 1 |
| CD4 (RPA-T4) | 1:120 | Brilliant Violet 785 | BioLegend (300554) | 1 |
| CD127 (A019D5) | 1:120 | Alexa Fluor 488 | BioLegend (351313) | 1 |
| CD45RA (HI100) | 1:120 | PE/Dazzle 594 | BioLegend (304146) | 1 |
| CD3 (UCHT1) | 1:120 | Brilliant Violet 605 | BioLegend (300460) | 2 |
| CD56 (5.1H11) | 1:120 | PE/Dazzle 594 | BioLegend (362544) | 2 |
| CD3 (UCHT1) | 1:120 | Brilliant Violet 605 | BioLegend (300460) | 3 |
| CD8 (RPA-T8) | 1:120 | Brilliant Violet 785 | BioLegend (301046) | 3 |
| CD45RA (HI100) | 1:120 | PE/Dazzle 594 | BioLegend (304146) | 3 |
| CD25 (M-A251) | 1:120 | Brilliant Violet 421 | BioLegend (356114) | 4 |
| CD3 (UCHT1) | 1:120 | Brilliant Violet 605 | BioLegend (300460) | 4 |
| CD4 (RPA-T4) | 1:120 | Brilliant Violet 785 | BioLegend (300554) | 4 |
| CD127 (A019D5) | 1:120 | Alexa Fluor 488 | BioLegend (351313) | 4 |
| CD45RA (HI100) | 1:120 | PE/Dazzle 594 | BioLegend (304146) | 4 |
| CD56 (5.1H11) | 1:120 | PE/Cy7 | BioLegend (362510) | 4 |
| CD8 (RPA-T8) | 1:120 | Alexa Fluor 647 | BioLegend (301062) | 4 |
| Foxp3 (259D) | 1:50 | Alexa Fluor 488 | BioLegend (320212) | 5 |
| CD25 (M-A251) | 1:120 | Brilliant Violet 421 | BioLegend (356114) | 5 |
| CD4 (SK3) | 1:120 | Brilliant Violet 605 | BioLegend (344646) | 5 |
| CD45RA (HI100) | 1:120 | PE/Dazzle 594 | BioLegend (304146) | 5 |
| pSTAT5 (C71E5) | 1:120 | Alexa Fluor 647 | CST* (9365) | 5 |
| CD3 (UCHT1) | 1:120 | Brilliant Violet 605 | BioLegend (300460) | 6 |
| CD8 (RPA-T8) | 1:120 | Alexa Fluor 647 | BioLegend (301062) | 6 |
| CD56 (5.1H11) | 1:120 | Alexa Fluor 488 | BioLegend (362518) | 6 |
| pSTAT5 (D4737) | 1:120 | PE | CST* (14603) | 6 |

### IL-2 variants’ mutations and conjugations

**Table S2:**
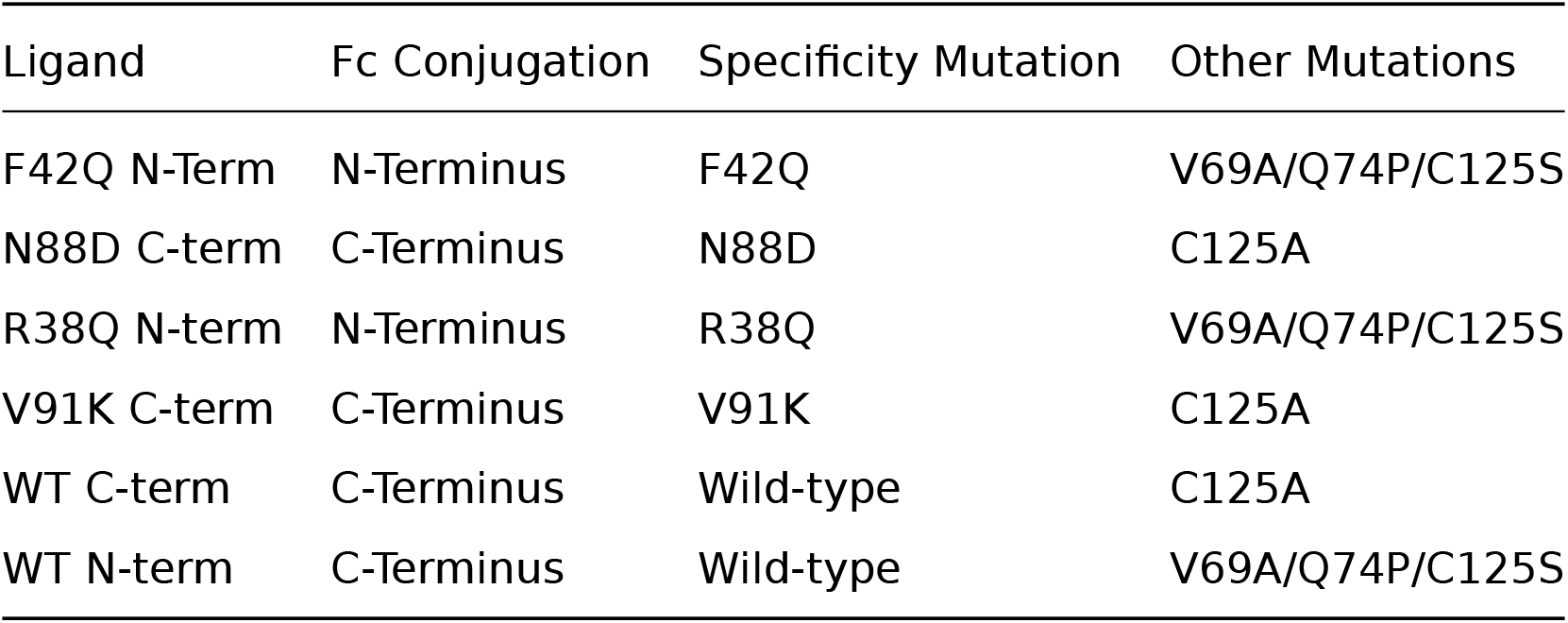
Modified IL-2 ligands and their respective mutations, and Fc conjugations.

| Ligand | Fc Conjugation | Specificity Mutation | Other Mutations |
| --- | --- | --- | --- |
| F42Q N-Term | N-Terminus | F42Q | V69A/Q74P/C125S |
| N88D C-term | C-Terminus | N88D | C125A |
| R38Q N-term | N-Terminus | R38Q | V69A/Q74P/C125S |
| V91K C-term | C-Terminus | V91K | C125A |
| WT C-term | C-Terminus | Wild-type | C125A |
| WT N-term | C-Terminus | Wild-type | V69A/Q74P/C125S |

### IL-2 variants’ IL-2Rβ/γ_c_ affinities

Data Table SD1: **IL-2Rβ/γ_c_ binding affinities of mutant and modified cytokines.** Data from the BLI studies for each IL-2 mutein.

**Figure S1:**
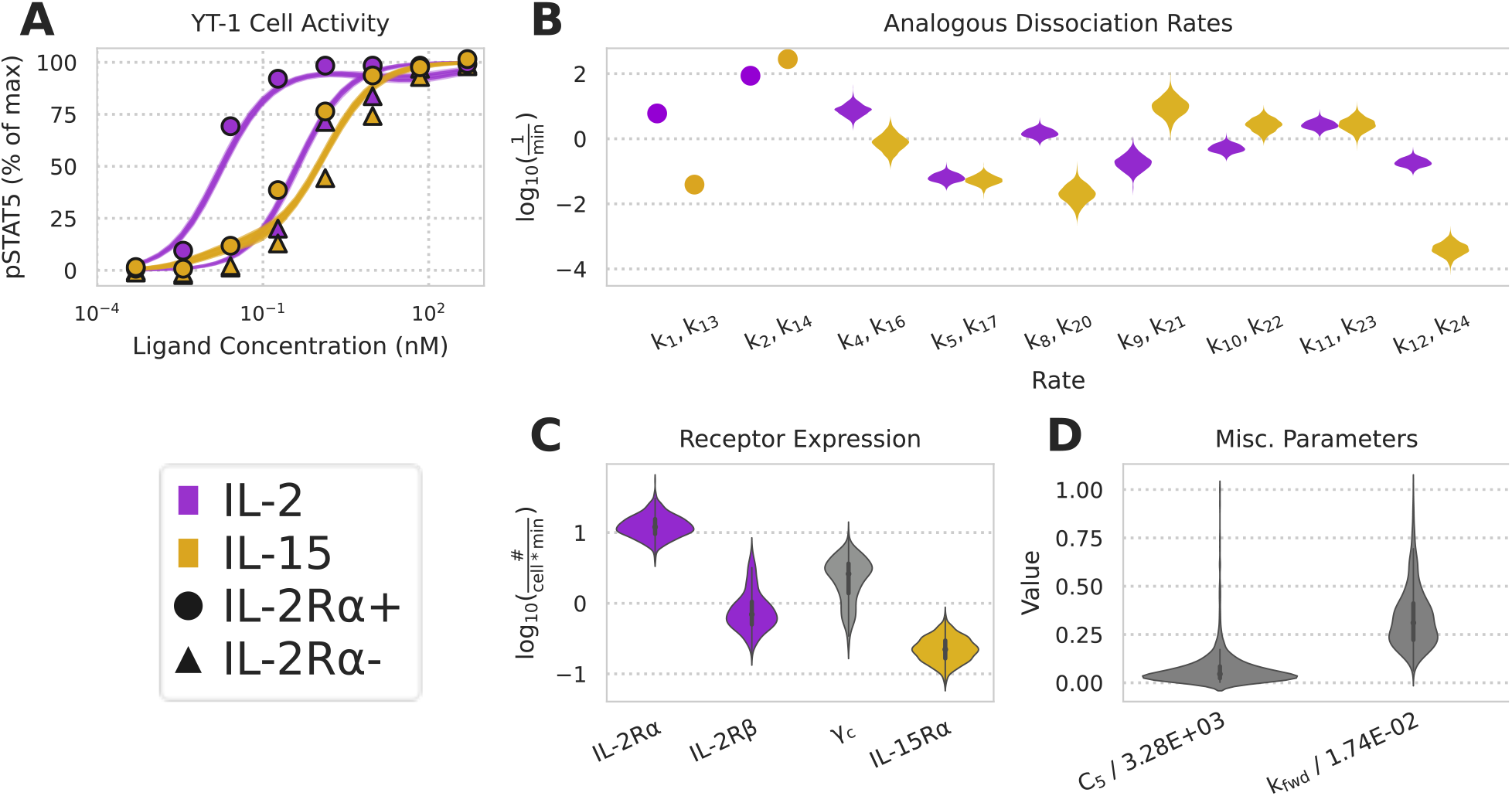
Model without trafficking fails to capture IL-2/-15 dose response. A) Model without trafficking fit to IL-2 and IL-15 pSTAT5 dose response data.^14^ This model was not fit to the surface IL-2Rβ measurements since no receptors were allowed to internalize from the cell surface (Fig. 1B-D). B) Posterior distributions of analogous reverse reaction rates for IL-2 and IL-15 in no-trafficking model. C) Posterior distributions for receptor surface abundance in no-trafficking model. D) Posterior distribution for the pSTAT5 activity scaling constant in no-trafficking model.

**Figure S2:**
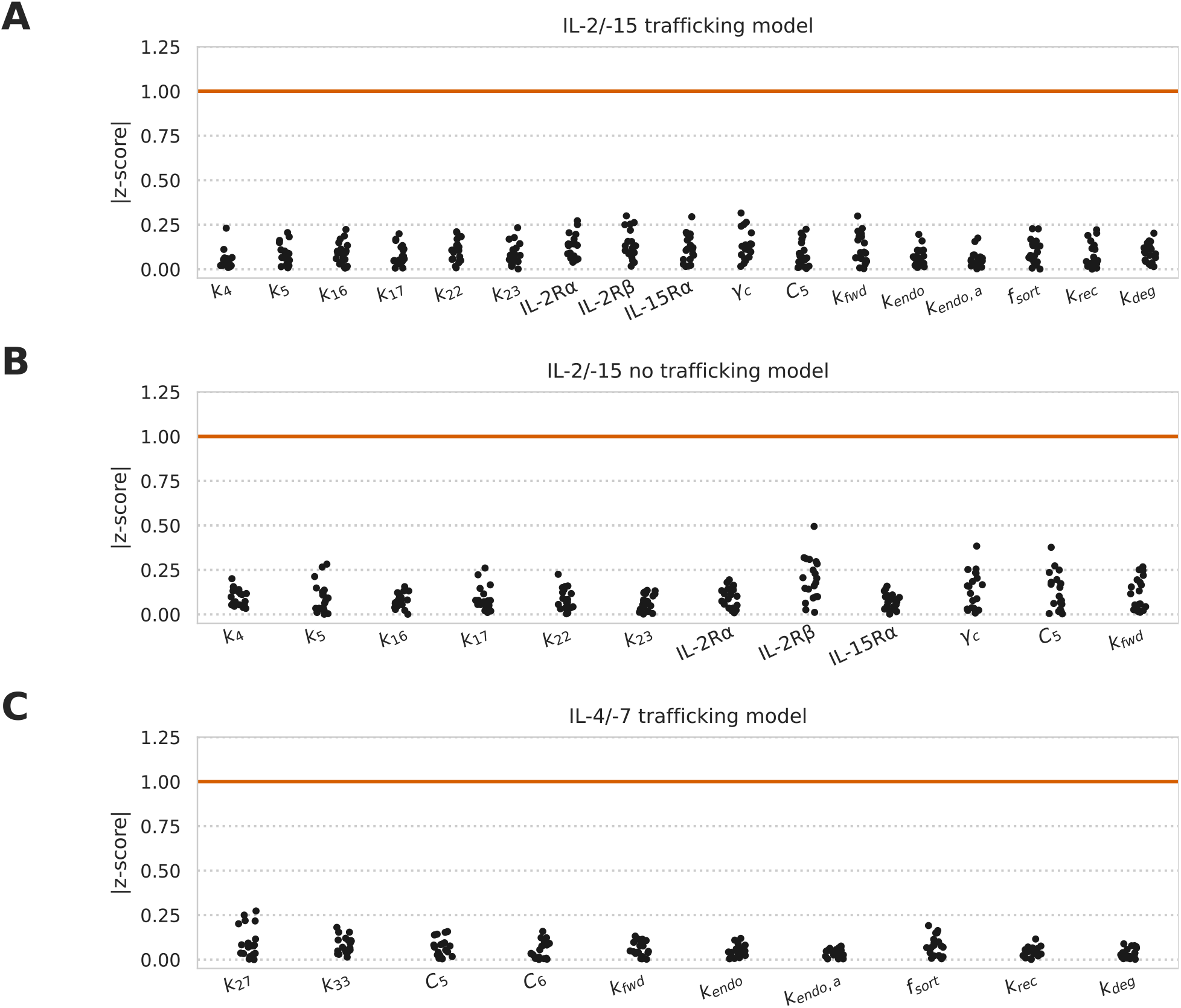
Geweke criterion scores for model fitting with and without trafficking. Geweke criterion z-scores in all subplots were calculated using 20 intervals in the first 10% and last 50% of MCMC chain. Scores of |z| < 1 imply fitting convergence. A-B) IL-2/-15 with and without trafficking. C) IL-4/-7 with trafficking (Fig. S1).

**Figure S3:**
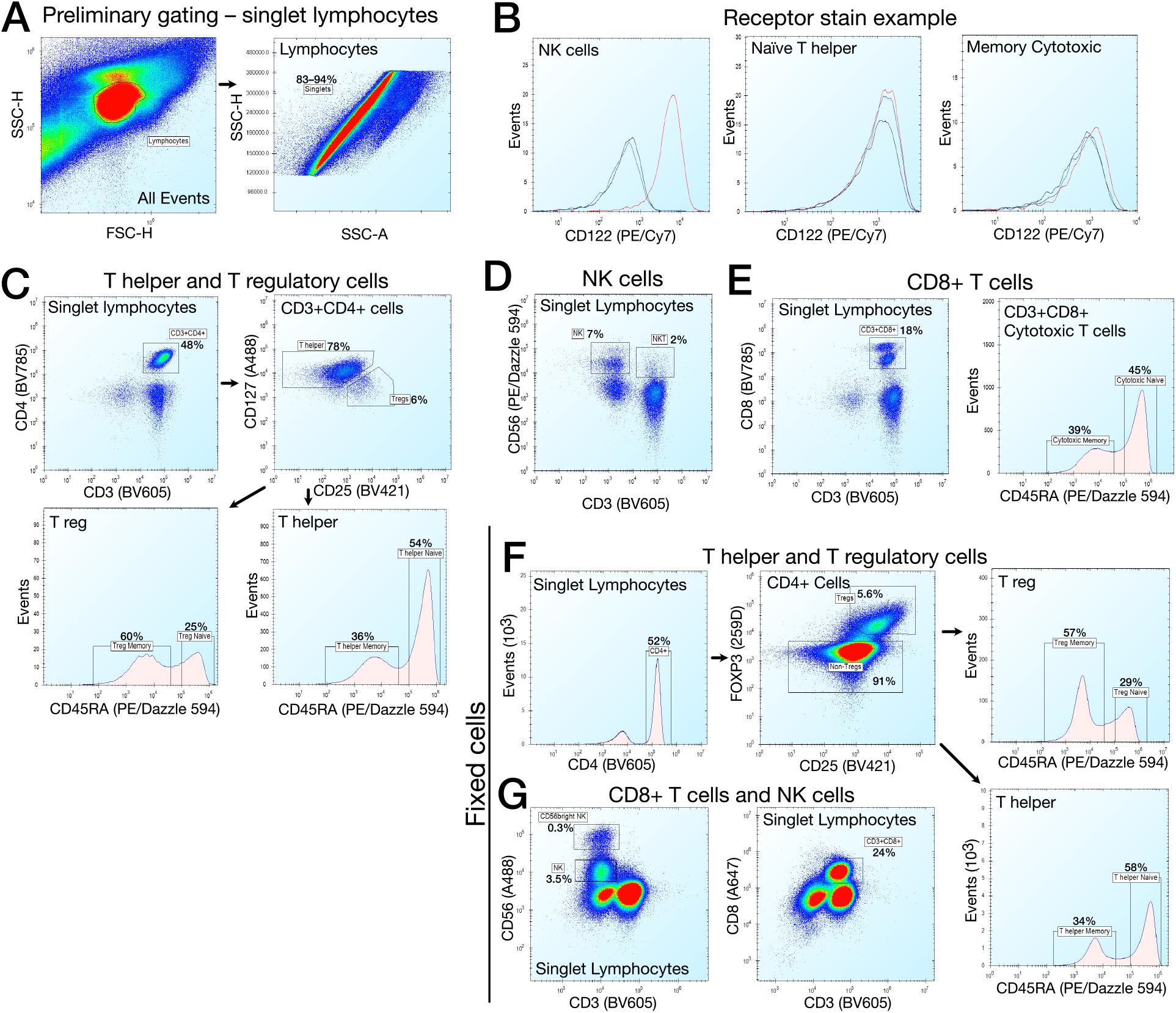
Receptor quantification and gating of PBMC-derived immune cell types. A) Preliminary gating for single lyphocytes. B) Example staining for CD122 (red), the corresponding isotype control (blue), and unstained cells (black). C) Gating for live T helper and T regulatory cells during receptor quantification. D) Live cell NK cell gating. E) Live cell CD8+ T cell gating. F) Gating for fixed T helper and T regulatory cells during pSTAT5 quantification. G) Fixed CD8+ T cell and NK cell gating.

**Figure S4:**
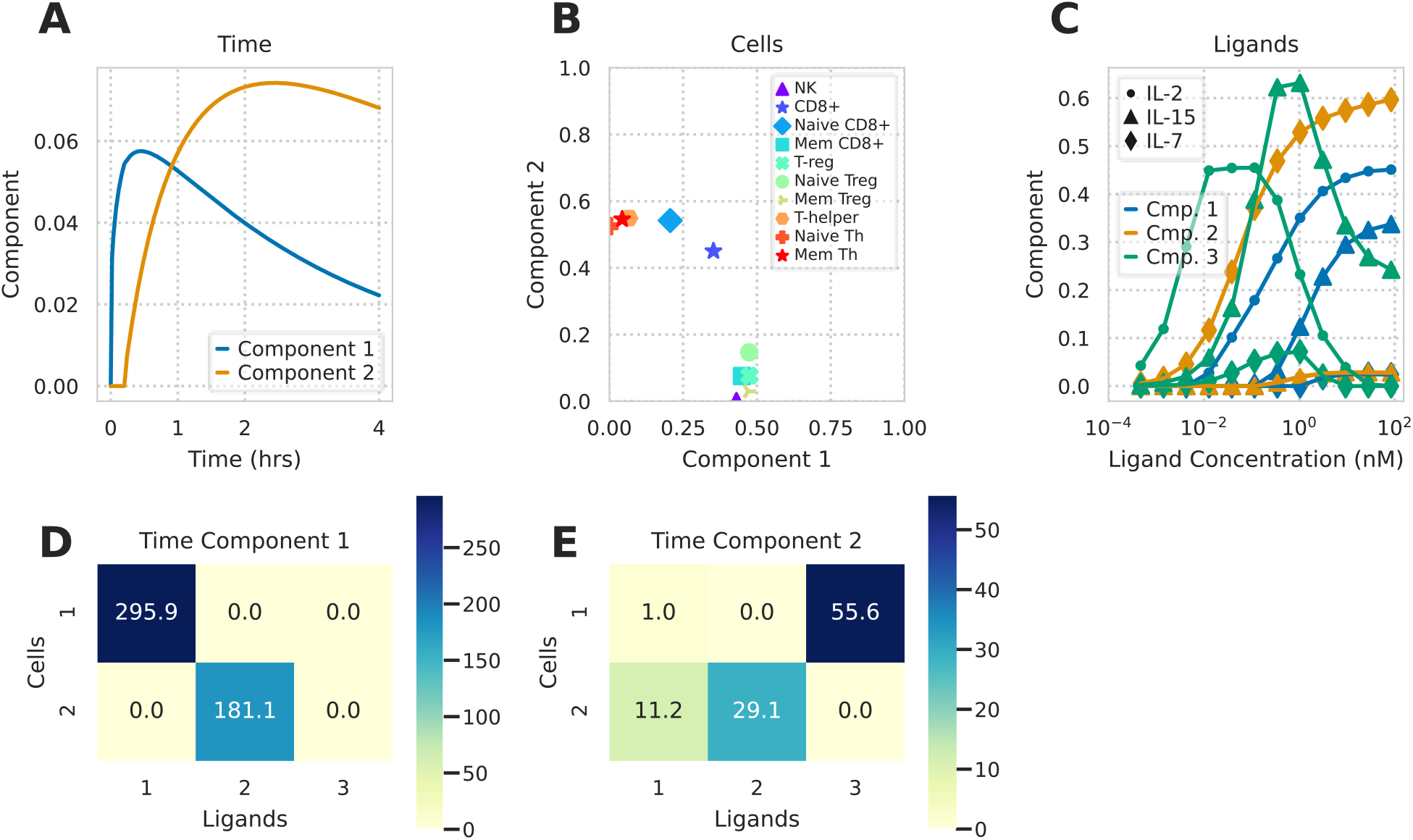
Tucker factorization of predicted immune cell type responses. A) Timepoint decomposition plot showing factorization component values against time after decomposing the tensor’s first dimension into 2 components. B) Decomposition plot along the second (cell) dimension after decomposing it to 2 components showing the ten cell type values along each component. C) Ligand decomposition plot along the tensor’s third dimension after decomposing it into 3 components. D–E) Slices of the Tucker core tensor corresponding to time component 1 (D) and 2 (E).

**Figure S5:**
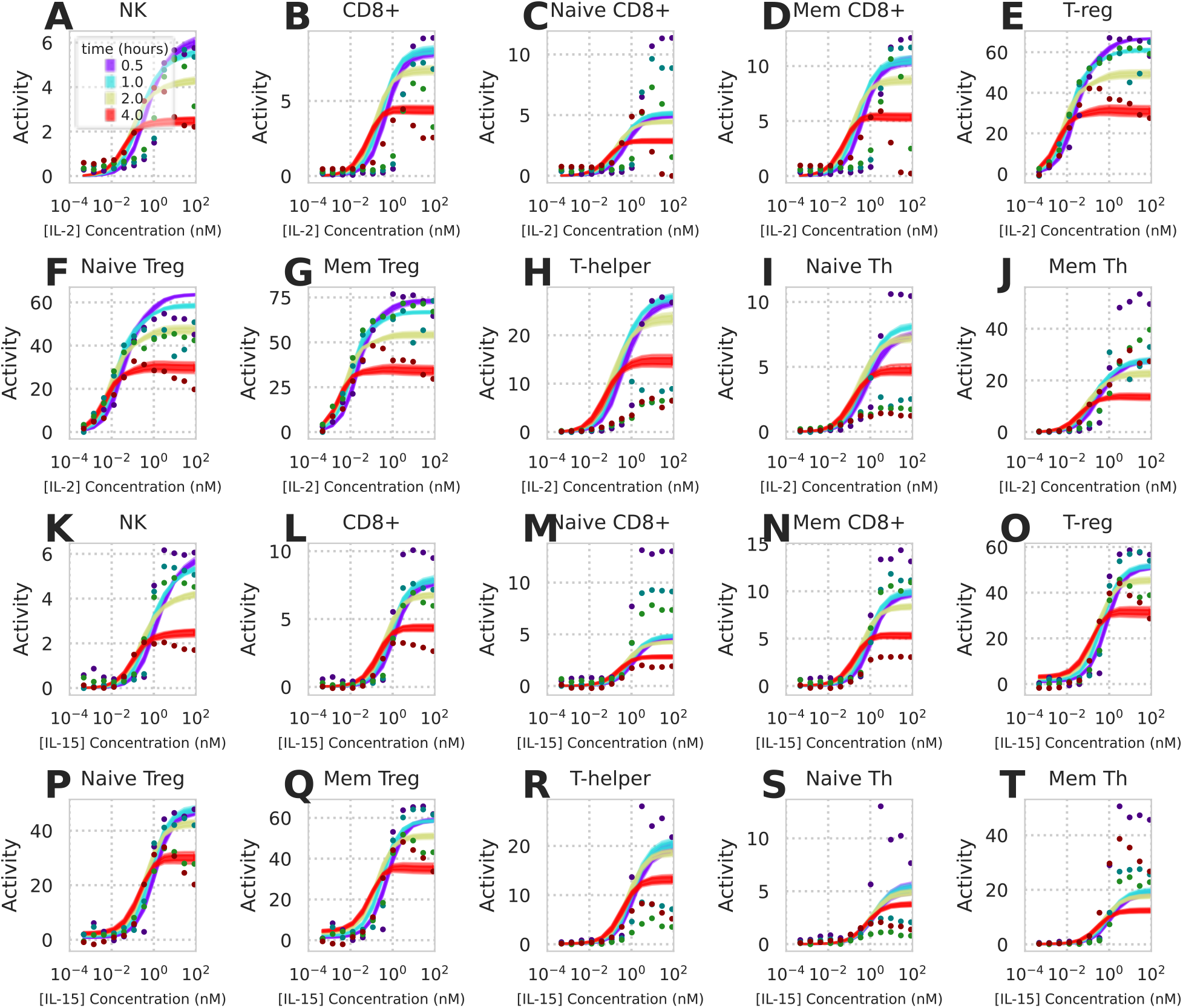
Full panel of predicted versus actual immune cell type responses. Dots represent experimental measurements and shaded regions represent 10-90% confidence interval for model predictions. Time of pSTAT5 activity measurement is denoted by color. All cell populations were stimulated with either IL-2 (A-J) or IL-15 (K-T).

**Figure S6:**
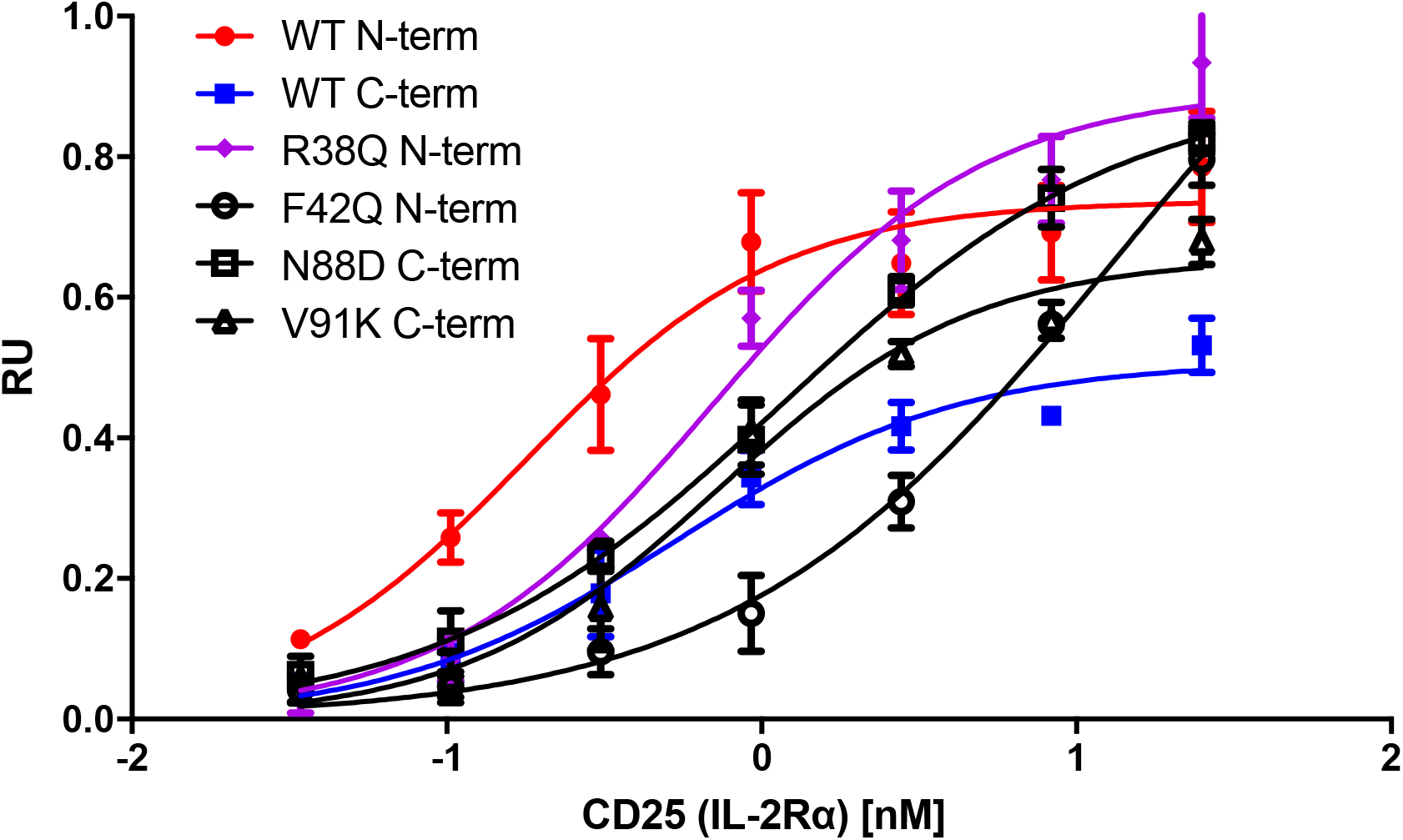
Cytokine affinity measurements to IL-2Rα. Binding is quantified in relative units.

**Figure S7:**
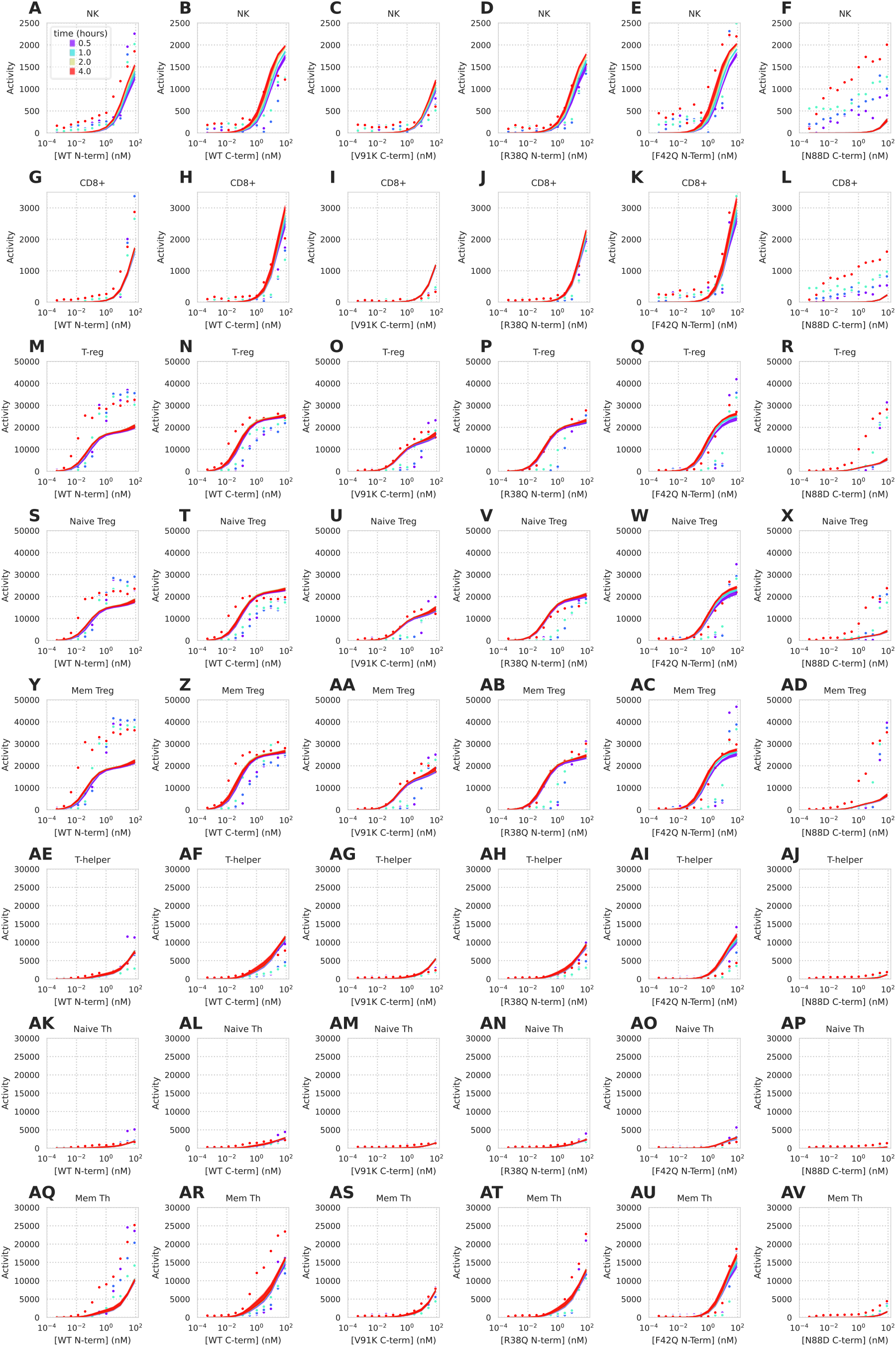
Full panel of predicted versus actual immune cell type responses to IL-2 muteins. Dots represent experimental measurements and shaded regions represent 10-90% confidence interval for model predictions. Time of pSTAT5 activity measurement is denoted by color. Cell populations were stimulated with IL-2 muteins of varying IL-2Rα and IL-2Rβ/γ_c_ binding affinities.

## References

1. Rochman, Y., Spolski, R. & Leonard, W. New insights into the regulation of T cells by γc family cytokines. Nature Reviews Immunology 9, 480–490 (2009).

2. Leonard, W. J., Lin, J.-X. & O’Shea, J. J. The γc family of cytokines: Basic biology to therapeutic ramifications. Immunity 50, 832–850 (2019).

3. Walsh, S. T. R. A biosensor study indicating that entropy, electrostatics, and receptor glycosylation drive the binding interaction between interleukin-7 and its receptor. Biochemistry 49, 8766–8778 (2010).

4. Amorosi, S. et al. The cellular amount of the common γ-chain influences spontaneous or induced cell proliferation. The Journal of Immunology 182, 3304–3309 (2009).

5. Vigliano, I. et al. Role of the common γ chain in cell cycle progression of human malignant cell lines. International Immunology 24, 159–167 (2012).

6. Wang, L. et al. Key role for IL-21 in experimental autoimmune uveitis. Proceedings of the National Academy of Sciences 108, 9542–9547 (2011).

7. Sharma, R. et al. Cutting edge: A regulatory t cell-dependent novel function of CD25 (IL-2Rα) controlling memory CD8+ T cell homeostasis. The Journal of Immunology 178, 1251–1255 (2007).

8. Sharfe, N., Dadi, H. K., Shahar, M. & Roifman, C. M. Human immune disorder arising from mutation of the α chain of the interleukin-2 receptor. Proceedings of the National Academy of Sciences 94, 3168–3171 (1997).

9. Horak, I. Immunodeficiency in il-2-knockout mice. Clin Immunol Immunopathol 76, S172–3

10. Human il2ra null mutation mediates immunodeficiency with lymphoproliferation and autoimmunity. Clinical Immunology 146, 248–261 (2013).

11. Stephanie R. Pulliam, S. E. A., Roman V. Uzhachenko & Shanker, A. Common gamma chain cytokines in combinatorial immune strategies against cancer. Immunology letters 169, 61–72 (2015).

12. Bentebibel, S.-E. et al. A first-in-human study and biomarker analysis of NKTR-214, a novel IL-2-receptor beta/gamma (βγ)-biased cytokine, in patients with advanced or metastatic solid tumors. Cancer Discovery (2019). doi:10.1158/2159-8290.CD-18-1495

13. Zhu, E. F. et al. Synergistic innate and adaptive immune response to combination immunotherapy with anti-tumor antigen antibodies and extended serum half-life il-2. Cancer Cell 27, 489–501

14. Ring, A. M. et al. Mechanistic and structural insight into the functional dichotomy between IL-2 and IL-15. Nature Immunology 13, 1187–1195 (2012).

15. Cotari, J. W., Voisinne, G., Dar, O. E., Karabacak, V. & Altan-Bonnet, G. Cell-to-cell variability analysis dissects the plasticity of signaling of common γ chain cytokines in T cells. Science Signaling 6, ra17–ra17 (2013).

16. Krieg, C., Letourneau, S., Pantaleo, G. & Boyman, O. Improved IL-2 immunotherapy by selective stimulation of IL-2 receptors on lymphocytes and endothelial cells. Proc Natl Acad Sci U S A 107, 11906–11911

17. Konrad, M. W. et al. Pharmacokinetics of recombinant interleukin 2 in humans. Cancer Research 50, 2009–2017 (1990).

18. Bernett, M. J. et al. Abstract 1595: IL-15/IL-15Rα heterodimeric Fc-fusions with extended half-lives. Cancer Research 77, 1595–1595 (2017).

19. Donohue, J. H. & Rosenberg, S. A. The fate of interleukin-2 after in vivo administration. The Journal of Immunology 130, 2203–2208 (1983).

20. William G. Berndt, K. A. S., David Z. Chang & Ciardelli, T. L. Mutagenic analysis of a receptor contact site on interleukin-2: Preparation of an IL-2 analog with increased potency. Biochemistry 33, 6571–6577

21. Collins, L. et al. Identification of specific residues of human interleukin 2 that affect binding to the 70-kDa subunit (p70) of the interleukin 2 receptor. Proceedings of the National Academy of Sciences 85, 7709–7713 (1988).

22. Aron M. Levin, A. M. R., Darren L. Bates. Exploiting a natural conformational switch to engineer an interleukin-2 ‘superkine’. Nature 484, 529–533 (2012).

23. Bell, C. J. M. et al. Sustained *in vivo* signaling by long-lived IL-2 induces prolonged increases of regulatory T cells. Journal of Autoimmunity 56, 66–80 (2015).

24. Peterson, L. B. et al. A long-lived IL-2 mutein that selectively activates and expands regulatory T cells as a therapy for autoimmune disease. Journal of Autoimmunity 95, 1–14 (2018).

25. Burke, M. A. et al. Modeling the proliferative response of T cells to IL-2 and IL-4. Cellular Immunology 178, 42–52 (1997).

26. Feinerman, O. et al. Single-cell quantification of IL-2 response by effector and regulatory T cells reveals critical plasticity in immune response. Molecular Systems Biology 6, (2010).

27. Gonnord, P. et al. A hierarchy of affinities between cytokine receptors and the common gamma chain leads to pathway cross-talk. Science Signaling 11, (2018).

28. Duprez, V., Cornet, V. & Dautry-Varsat, A. Down-regulation of high affinity interleukin 2 receptors in a human tumor T cell line. Interleukin 2 increases the rate of surface receptor decay. Journal of Biological Chemistry 263, 12860–12865 (1988).

29. Lamaze, C. et al. Interleukin 2 receptors and detergent-resistant membrane domains define a clathrin-independent endocytic pathway. Molecular Cell 7, 661–671 (2001).

30. Eubelen, M. et al. A molecular mechanism for Wnt ligand-specific signaling. Science 361, (2018).

31. Li, P. et al. Morphogen gradient reconstitution reveals Hedgehog pathway design principles. Science 360, 543–548 (2018).

32. Antebi, Y. E., Nandagopal, N. & Elowitz, M. B. An operational view of intercellular signaling pathways. Current Opinion in Systems Biology 1, 16–24 (2017).

33. Antebi, Y. E. et al. Combinatorial signal perception in the BMP pathway. Cell 170, 1184–1196

34. Fallon, E. M. & Lauffenburger, D. A. Computational model for effects of lig-and/receptor binding properties on interleukin-2 trafficking dynamics and T cell proliferation response. Biotechnology Progress 16, 905–916

35. Fallon, E. M., Liparoto, S. F., Lee, K. J., Ciardelli, T. L. & Lauffenburger, D. A. Increased endosomal sorting of ligand to recycling enhances potency of an interleukin-2 analog. Journal of Biological Chemistry 275, 6790–6797 (2000).

36. Basquin, C. et al. The signalling factor PI3K is a specific regulator of the clathrin-independent dynamin-dependent endocytosis of IL-2 receptors. Journal of Cell Science 126, 1099–1108 (2013).

37. Volkó, J. et al. IL-2 receptors preassemble and signal in the er/golgi causing resistance to antiproliferative anti-il-2Rα therapies. Proceedings of the National Academy of Sciences 116, 21120–21130 (2019).

38. Mitra, S. et al. Interleukin-2 activity can be fine tuned with engineered receptor signaling clamps. Immunity 42, 826–838 (2015).

39. Spangler, J. B. et al. Antibodies to interleukin-2 elicit selective T cell subset potentiation through distinct conformational mechanisms. Immunity 42, 815–825 (2015).

40. Hassan, J. & Reen, D. J. IL-7 promotes the survival and maturation but not differentiation of human post-thymic CD4+ T cells. European journal of immunology 28, 3057–3065 (1998).

41. Tucker, L. R. Some mathematical notes on three-mode factor analysis. Psychometrika 31, 279–311 (1966).

42. Junghans, R. P. & Waldmann, T. A. Metabolism of tac (IL2Ralpha): Physiology of cell surface shedding and renal catabolism, and suppression of catabolism by antibody binding. Journal of Experimental Medicine 183, 1587–1602 (1996).

43. Kuwabara, T., Kasai, H. & Kondo, M. Acetylation modulates IL-2 receptor signaling in T cells. Journal of Immunology 197, 4334–4343 (2016).

44. Casim A. Sarkar, T. H., Ky Lowenhaupt. Rational cytokine design for increased lifetime and enhanced potency using pH-activated ‘histidine switching’. Nature Biotechnology 20, 908–913 (2002).

45. Haugh, J. M. Mathematical model of human growth hormone (hGH)-stimulated cell proliferation explains the efficacy of hGH variants as receptor agonists or antagonists. Biotechnology Progress 20, 1337–1344

46. Meyer, A. S., Zweemer, A. J. & Lauffenburger, D. A. The AXL receptor is a sensor of ligand spatial heterogeneity. Cell Systems 1, 25–36 (2015).

47. Leon, K., Garcia-Martinez, K. & Carmenate, T. Mathematical models of the impact of IL-2 modulation therapies on T cell dynamics. Frontiers in Immunology 4, 439 (2013).

48. Komorowski, M. & Tawfik, D. S. The limited information capacity of cross-reactive sensors drives the evolutionary expansion of signaling. Cell Systems 8, 76–85.e6 (2019).

49. Robinett, R. A. et al. Dissecting fcγr regulation through a multivalent binding model. Cell Systems (2018). doi:10.1016/j.cels.2018.05.018

50. Voss, S. D., Leary, T. P., Sondel, P. M. & Robb, R. J. Identification of a direct interaction between interleukin 2 and the p64 interleukin 2 receptor gamma chain. Proceedings of the National Academy of Sciences 90, 2428–2432 (1993).

51. Walsh, S. T. R. Structural insights into the common γ-chain family of cytokines and receptors from the interleukin-7 pathway. Immunological Reviews 250, 303–316

52. Renauld, J. C. et al. Expression cloning of the murine and human interleukin 9 receptor cDNAs. Proceedings of the National Academy of Sciences 89, 5690–5694 (1992).

53. Rickert, M., J Boulanger, M., Goriatcheva, N. & Christopher Garcia, K. Compensatory energetic mechanisms mediating the assembly of signaling complexes between interleukin-2 and its α, β, and γc receptors. Journal of molecular biology 339, 1115–28 (2004).

54. Mortier, E. et al. Soluble interleukin-15 receptor α (IL-15Rα)-sushi as a selective and potent agonist of IL-15 action through IL-15Rβ/γ: Hyperagonist IL-15oIL-15Rα fusion proteins. Journal of Biological Chemistry 281, 1612–1619 (2006).

55. Dubois, S., Mariner, J., Waldmann, T. A. & Tagaya, Y. IL-15Rα recycles and presents IL-15 in *trans* to neighboring cells. Immunity 17, 537–547 (2002).

56. Hindmarsh, A. C. et al. SUNDIALS: Suite of nonlinear and differential/algebraic equation solvers. ACM Transactions on Mathematical Software (TOMS) 31, 363–396 (2005).

57. Cao, Y., Li, S. & Petzold, L. Adjoint sensitivity analysis for differential-algebraic equations: Algorithms and software. Journal of Computational and Applied Mathematics 149, 171–191 (2002).

58. Hogan, R. J. Adept 2.0: a combined automatic differentiation and array library for C++. (2017). doi:10.5281/zenodo.1004730

59. Geweke, J. Evaluating the accuracy of sampling-based approaches to the calculation of posterior moments. in Bayesian statistics 4 169–193 (University Press, 1992).

60. Kossaifi, J., Panagakis, Y. & Pantic, M. TensorLy: Tensor learning in Python. CoRR abs/1610.09555, (2016).

61. Ishino, T. et al. Engineering a monomeric Fc domain modality by N-glycosylation for the half-life extension of biotherapeutics. Journal of Biological Chemistry 22, 473– 482 (2013).

